# Early spatial and contextual coding deficits in hippocampal CA1 precede performance decline in an Alzheimer’s disease model

**DOI:** 10.1101/2025.02.05.636661

**Authors:** Yimei Li, Mary Ann Go, Hualong Zhang, Robertas Aleksynas, Juliana Kim, Mingyang Gao, Seigfred Prado, Magdalena Sastre, Simon R Schultz

## Abstract

Alzheimer’s disease (AD) is characterized by progressive memory decline, yet how hippocampal representations change at early AD stages remains largely unknown. In this study, we utilized a real-world head-fixed spatial alternation task combined with in vivo two-photon imaging to investigate hippocampal CA1 representations in 5xFAD mice. While 7-9 month old 5xFAD mice showed significant deficits in task learning, 2-4 month old mice performed normally, enabling assessment of hippocampal coding prior to behavioral decline. At this early stage, CA1 neurons exhibited intact global spatial encoding but demonstrated impaired task-related place cell representations. Contextual representations revealed by trajectory-dependent coding were weakened at both the population and single cell levels, accompanied by a notable shift from predominantly retrospective to more prospective encoding during behavioral choices. During task learning, longitudinally unstable place cells showed greater impairments in spatial coding, whereas stable place cells remained largely preserved in early-stage AD mice. Additionally, chemogenetic activation of basal forebrain cholinergic neurons increased spatial information and within-session stability, while promoting trajectory-dependent coding in these early-stage 5xFAD mice. Together, these findings reveal selective early impairments in hippocampal task-related spatial and contextual representations in AD that precede overt deficits in spatial alternation performance and remains amenable to cholinergic modulation.

## Introduction

Alzheimer’s disease (AD) is the most prevalent cause of dementia in older adults, particularly affecting individuals over 65 years of age ^1^. A central feature of AD pathology is the amyloidogenic processing of amyloid precursor protein (APP), which produces amyloid-β (Aβ) and triggers downstream pathophysiological cascades leading to synaptic dysfunction and neuronal loss ^2, 3, 4, 5, 6^. Among the cognitive impairments associated with AD, spatial navigation deficits are especially prominent and often emerge as early as the mild cognitive impairment stage ^7, 8^. More than 60% of AD patients experience spatial disorientation and wandering behavior ^9^. These deficits are closely linked to hippocampal dysfunction. Hippocampal CA1 neurons, crucial for forming spatial and episodic memories, exhibit abnormalities in morphology and synaptic plasticity in both human patients with AD and mouse models ^10, 11, 12^.

Hippocampal place cells are essential for supporting spatial navigation and memory formation, and accumulating evidence indicates that their function is disrupted in AD. Impairments in place cell firing have been reported across several AD mouse models, including APP transgenic and APP knock-in lines. For example, 11-16 month old Tg2576 and APP mice show degraded place fields and reduced spatial information, typically in conjunction with behavioral deficits ^13, 14^. In 3xTg AD mice, reduced spatial information has been reported in young 3xTg mice during open-field or linear track running, with reduced theta- and slow-gamma phase locking ^15, 16^. In 5xFAD mice, impaired synaptic efficacy and weakened sharp wave-ripples have been observed during virtual reality spatial tasks at 11–14 months of age^17^. Overall, these findings demonstrate place cell dysfunction in AD, although much of the evidence comes from later disease stages or from exploratory paradigms rather than behaviorally demanding tasks.

Recent longitudinal calcium-imaging work showed that CA1 spatial representations in 5xFAD mice become unreliable at an early disease stage, before object-location memory deficits emerge. In 4–5 month old 5xFAD mice, spatial coding abnormalities were observed during open-field exploration and linear-track running, whereas object-location task performance remained intact ^18^. However, how early AD affects task-related spatial and contextual representations within the same ongoing goal-directed task remains unclear.

In contrast to place cells, which encode spatial location, splitter cells encode trajectory-dependent contextual information, firing differently on a common path depending on past choices or future decisions. First identified in continuous alternation tasks in rats ^19, 20^, splitter cells integrate retrospective and prospective information within single CA1 neurons and are thought to support flexible navigation ^21^. Despite their importance, little is known about how splitter cell activity is affected in AD. Given that Aβ-related synaptic changes impair hippocampal learning ^22^ and that the emergence of splitter cell activity can be driven by rapid synaptic plasticity ^23^, trajectory-dependent contextual coding may be particularly vulnerable early in the disease.

Here, we developed a real-world head-fixed continuous Phi-maze alternation task that imposes goal-directed spatial learning demands comparable to those in freely moving paradigms. We combined this task with two-photon calcium imaging in 5xFAD mice, which begin to accumulate amyloid pathology around 2-4 months of age ^24, 25^, allowing us to examine spatial and trajectory-dependent contextual representations within the same behavioral framework. We examined early CA1 coding abnormalities at multiple levels, including task-related spatial representations, the vulnerability of distinct place cell populations, trajectory-dependent contextual coding, and the representation of previous and upcoming trajectory information. Given the established role of basal forebrain cholinergic input in modulating hippocampal synaptic plasticity ^26^, we then tested whether cholinergic activation could modulate these early coding abnormalities in 5xFAD mice.

## Results

### 1. Spatial learning and global CA1 spatial coding are preserved in young 5xFAD mice but impaired with age

We developed a novel behavioral paradigm to assess spatial memory in head-fixed mice, in which animals performed a continuous spatial alternation task within a real-world, air-lifted Phi-maze (Figure 1a, left). Mice received a 4 μl of liquid reward only when they entered the arm opposite to that visited in the previous trial (Figure 1a, right). Water-restricted mice expressing jGCaMP7s in CA1 pyramidal neurons of the dorsal hippocampus were trained to perform the task while calcium activity was simultaneously recorded using two-photon microscope (Figure 1b).

**Figure 1.**
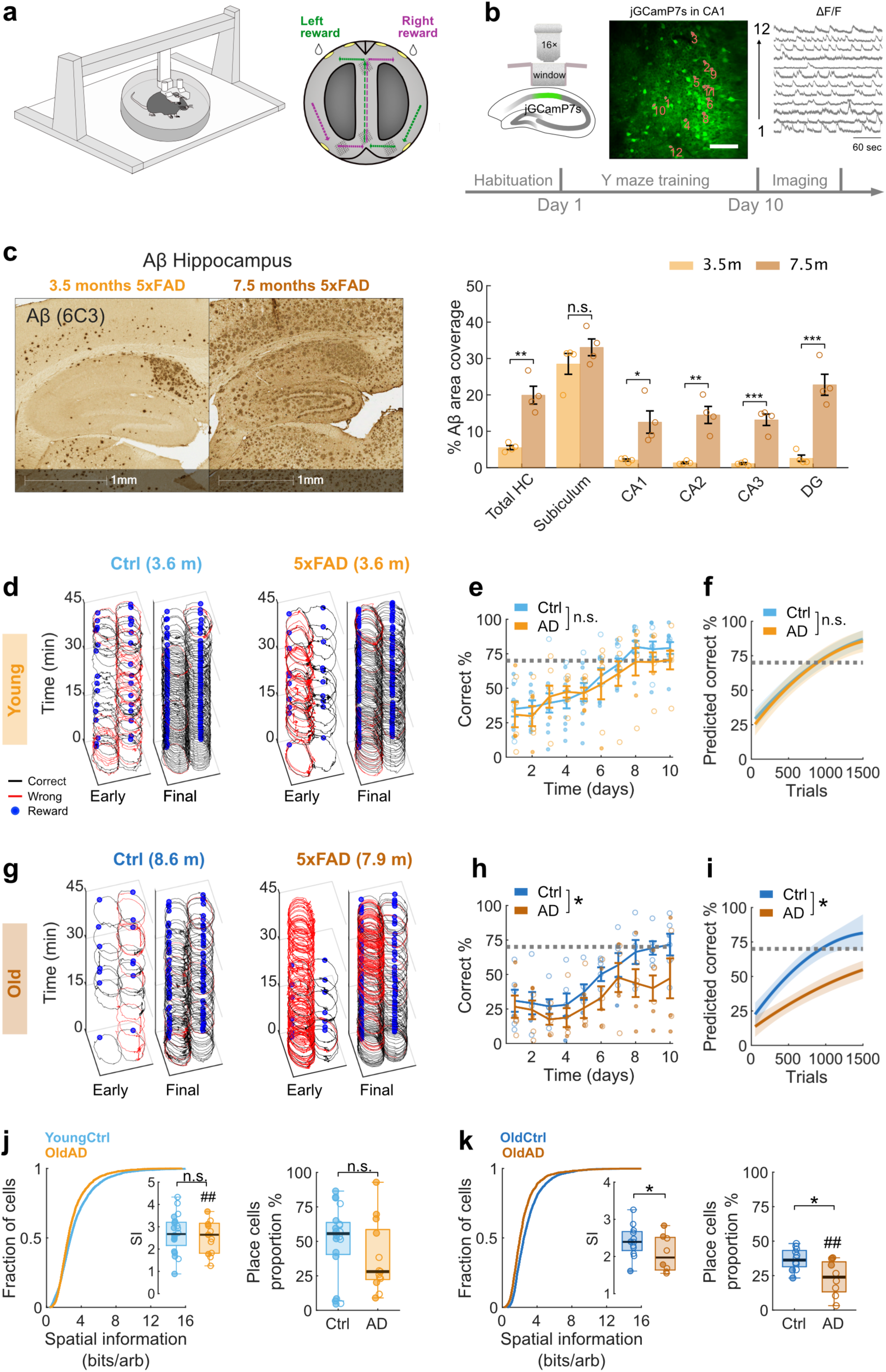
Age-dependent behavioral deficits and global CA1 spatial coding in 5xFAD mice during the Phi-maze alternation task. **a.** Left, schematic of a head-fixed mouse navigating an air-lifted Phi-maze. Right, bird’s-eye view of the Phi-maze. Correct left-to-right and right-to-left trials are indicated in purple and green, respectively. **b.** Schematic of in vivo calcium imaging from CA1 pyramidal neurons expressing jGCaMP7s. Twelve example regions of interest (ROIs) are overlaid on the imaging field of view, with corresponding calcium ΔF/F traces shown on the right. Scale bar, 100 µm. **c.** Aβ deposition in 5xFAD mice at 3.5 and 7.5 months of age. Left, representative hippocampal images from 30-µm sagittal sections stained with the pan-Aβ antibody MOAB-2 (clone 6C3) and visualised by DAB. Right, quantification of Aβ positive area in hippocampal subregions and the total hippocampus. **d.** Representative behavioral trajectories during early and final training days in 2-4-month old mice. Group median age is shown above each group. Correct trials are shown in black, error trials in red and reward locations in blue. **e.** Learning curves showing the percentage of correct alternation trials over training days in 2-4-month-old mice. The horizontal dotted line indicates the 70% performance criterion. Ctrl, n = 10 mice; AD, n = 7 mice. Genotype × day interaction, linear mixed-effects model (LMM). **f.** Percentage of correct trials across consecutive 50-trial blocks in 2-4-month-old mice. Curves show quadratic mixed-effects model fits, with shaded areas indicating 95% confidence intervals. Genotype × block interaction, LMM. **g-i.** Same as in panels **d-f**, but for older 5xFAD mice (> 7 months). Ctrl, n = 6 mice; AD, n = 6 mice. **j.** Left, cumulative distribution of Skaggs’ spatial information for all detected CA1 neurons, based on two-dimensional spatial maps from 2-4 month old mice. LMM test. Inset, session level medians. Right, proportion of neurons classified as place cells per session. Two-sided Wilcoxon rank-sum tests. Ctrl, n = 4,397 cells from 20 imaging sessions in 10 mice; AD, n = 3,073 cells from 13 sessions in 7 mice. **k.**Same as in panel **j**, but for old 5xFAD mice. Ctrl, n = 2,168 cells from 12 imaging sessions in 6 mice; AD, n = 1,477 cells from 8 sessions in 6 mice. Asterisks denote significant genotype effects, whereas hash symbols denote significant genotype × sex interactions, with lower values in female 5xFAD mice. n.s., not significant; *P < 0.05; **P < 0.01; ***P < 0.001. Mouse and cell information are provided in Supplementary Table 1.

Consistent with previous reports^27^, Methoxy-X04 labeling showed widespread Aβ deposition throughout the brain in 8 months old 5xFAD mice, and 6C3 staining further revealed age-dependent Aβ accumulation in the hippocampus of 5xFAD mice (Figure 1c and Supplementary Figure 1). The percentage of Aβ positive area increased from 3.5 to 7.5 months in the total hippocampus (5.53 ± 0.57% vs 19.92 ± 2.45%, p = 0.001; Figure 1c) and CA1 (2.11 ± 0.28% vs 12.54 ± 3.06%, p = 0.015). Similar increases were observed in CA2, CA3 and DG. We therefore compared young mice around 3 months of age (Ctrl, median age 3.6 months; 5xFAD, median age 3.6 months) with mice older than 7 months (Ctrl, median age 8.6 months; 5xFAD, median age 7.9 months) to examine task performance. Animal details are provided in Supplementary Table 1.

In young mice, 5xFAD mice performed comparably to age matched controls (Genotype × Day, β = -1.29, 95% CI [-2.63, 0.44], p = 0.16, linear mixed-effects model (LMM); Figure 1d,e). In contrast, old 5xFAD mice exhibited pronounced learning impairments (Genotype × Day, β = -2.56, 95% CI [-4.57, -0.54], p = 0.013; Figure 1g,h). These behavioral effects did not show a significant Genotype × Sex × Day interaction in either age group (young, p = 0.769; old, p = 0.064), indicating that the learning phenotype was not statistically sex dependent. Notably, old 5xFAD mice also showed greater variability in the number of trials completed per day, potentially affecting learning opportunities (Supplementary Figure 2a-f). To account for this, trials from all sessions were grouped into consecutive 50-trial blocks, and the percentage of correct trials was calculated per block (Supplementary Figure 2g,h). This analysis yielded results consistent with day-wise performance: young 5xFAD mice displayed no significant behavioral deficits (Genotype × Block, β = 0.35, 95% CI [-0.70, 1.40], p = 0.51; Figure 1f), whereas old 5xFAD mice showed a significant impairment (Genotype × Block, β = -1.85, 95% CI [-3.63, -0.06], p = 0.043; Figure 1i).

Furthermore, the proportion of old 5xFAD mice reaching the task acquisition criterion was significantly reduced compared with controls, whereas young 5xFAD mice showed normal acquisition rates (Supplementary Figure 2g,h). These findings indicate that spatial learning deficits are evident in old 5xFAD mice, consistent with previous studies ^25^, while while young 5xFAD mice retain intact task performance.

To evaluate overall CA1 spatial coding, we quantified global spatial information, defined as the Skaggs’ spatial information content computed from two-dimensional spatial activity maps across all trajectories within the imaging session. In young mice, CA1 neurons from 5xFAD mice exhibited spatial information levels comparable to controls (β = -0.12, 95% CI [-0.89, 0.66], p = 0.77; LMM; Figure 1j, left), with a significant Genotype × Sex interaction (p = 0.003). The proportion of neurons meeting the place cell criterion also showed a non-significant reduction in young 5xFAD mice (r_rb_ = -0.25, p = 0.25; Figure 1j, right), with no significant genotype × sex interaction (p = 0.15). In contrast, old 5xFAD mice displayed significant impairments in spatial coding, as reflected by reduced global spatial information (β = -0.54, 95% CI [-1.00, -0.09], p = 0.019; LMM; Figure 1k, left), with no significant genotype × sex interaction (p = 0.30). Old 5xFAD mice also showed a lower proportion of place cells (r_rb_ = -0.67, p = 0.015; Figure 1k, right), accompanied by a significant genotype × sex interaction (p = 0.002). Together, these results demonstrate that young 5xFAD mice retain intact spatial learning and broadly preserved global CA1 spatial coding during the Phi-maze alternation task, whereas both behavioral performance and neuronal spatial representations are markedly impaired in old 5xFAD mice.

### 2. Task-related CA1 place-cell coding is impaired in young 5xFAD mice

Understanding the early stage of AD is crucial for identifying initial circuit level disruptions and informing early intervention ^28, 29^. Based on our behavioral, pathological and neuronal observations (Figure 1c-k), together with previous reports of impaired hippocampal long-term potentiation and the emergence of detectable hippocampal-dependent behavioral deficits with disease progression in 5xFAD mice ^27, 30, 31^, we focused on young 5xFAD mice to examine early alternations in hippocampal neural coding.

To assess task-related spatial representations, we linearised trajectories and generated one-dimensional spatial activity maps during the post acquisition phase, when mice consistently performed the alternation task. Analyses were restricted to correct trials, which most directly reflect task engagement. This analysis therefore assessed task-aligned spatial coding, rather than global spatial tuning across all trajectories.

We first examined CA1 place cell activity during correct trials (Figure 2a). Behavioral performance and locomotion during imaging sessions were comparable between young control and 5xFAD mice, indicating that any differences in neuronal coding were not driven by overt correct performance (r_rb_ = 0.008, p = 0.99) or locomotor changes (r_rb_ = - 0.21, p = 0.33; Figure 2b and r_rb_ = 0.11, p = 0.62; Supplementary Figure 3b). Somatic calcium transient decay kinetics were also comparable between young control and 5xFAD mice, arguing against genotype-dependent differences in calcium signal kinetics as an explanation for the early coding phenotype. In contrast, calcium transient decay was shorter in old 5xFAD mice than in age matched controls, suggesting that calcium signal kinetics may become altered at later disease stages (Supplementary Figure 4).

**Figure 2.**
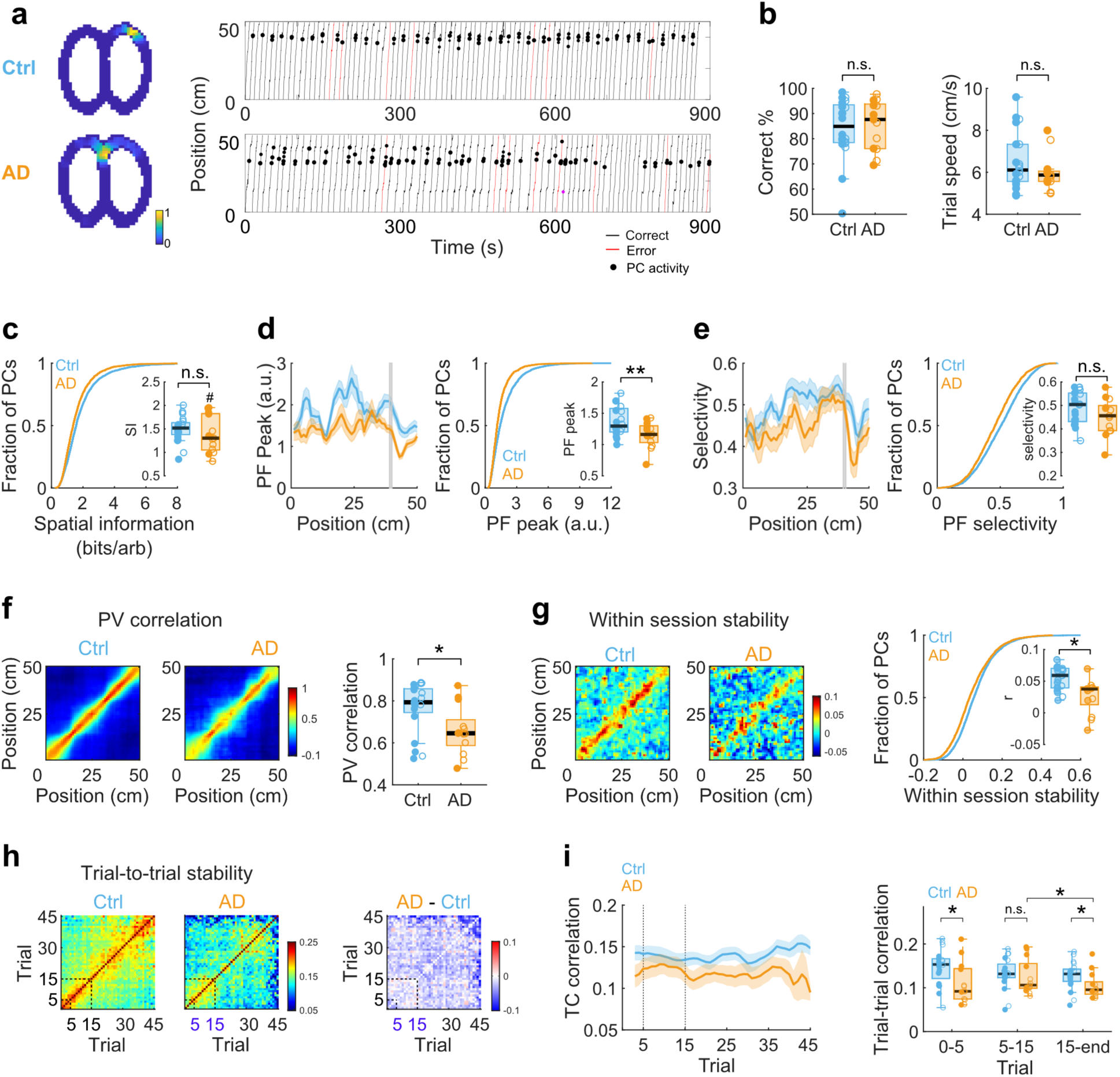
Impaired CA1 place cell activity during correct trials in young 5xFAD mice. **a.** Representative CA1 place cells recorded from young control and 5xFAD mice during the Phi-maze alternation task. Left, normalized two-dimensional spatial activity maps. Right, linearized behavioral trajectories and place cell activity across trials. Black lines indicate correct trials, red lines indicate error trials and black dots indicate place cell calcium events. **b.** Behavioral performance during imaging sessions. Left, percentage of correct alternation trials. Right, median of trial speeds across correct trials. **c.** Cumulative distribution of Skaggs’ spatial information for CA1 place cells during correct trials. Inset, session-level medians. **d.** Peak in-field activity of place fields. Left, in-field peak activity along the linearized trajectory. Shaded areas indicate mean ± s.e.m. across place fields. Right, cumulative distribution of in-field peak activity. Inset, session-level medians. **e.** Same as in panel **d**, but for in-field versus out-of-field selectivity, calculated as (in-field − out-of-field) / (in-field + out-of-field). **f.** Population vector correlations between odd and even correct trials across all detected CA1 neurons. Left, group averaged PV correlation matrices. Right, mean diagonal PV correlation for each imaging session. Two-sided Wilcoxon rank-sum tests. **g.** Within-session stability of place cell tuning. Left, representative odd-even trials tuning curve correlation matrices for control and 5xFAD mice. Right, cumulative distribution of mean within-session stability across place cells. Inset, session-level medians. **h.** Trial-to-trial stability of place cell tuning during correct trials. Left and middle, trial-by-trial tuning curve correlation matrices for the first 30 correct trials in control and 5xFAD mice. Right, difference map showing 5xFAD minus control correlations. **i.** Left, sliding-window analysis of trial-to-trial tuning stability across the session, tuning curve correlation plotted as a function of trial number. Right, tuning curve correlations for early trials (1-5), intermediate trials (5-15) and later trials (15 to end trial of the session). Two-sided Wilcoxon rank-sum tests. Ctrl, 4,397 recorded cells and 2,952 place cells from 20 imaging sessions in 10 mice; 5xFAD, 3,073 recorded cells and 1,791 place cells from 13 imaging sessions in 7 mice. Cell-level comparisons in panels c-e and g were performed using linear mixed-effects models (LMMs). Filled circles indicate male mice and open circles indicate female mice. Asterisks denote significant genotype effects; hash symbols denote significant Genotype × Sex interactions. n.s., not significant; *P < 0.05; **P < 0.01. Mouse and cell information are provided in Supplementary Table 1.

In 5xFAD mice, place cells showed a trend toward reduced spatial information relative to controls (β = -0.24, 95% CI [-0.48, 0.006], p = 0.056; Figure 2c), with a significant Genotype × Sex interaction (p = 0.01), indicating that the reduction apparent in females but not males. Place field fidelity was then assessed using peak in-field activity and in-field versus out-of-field selectivity. Peak in-field activity was significantly lower in 5xFAD mice (β = -0.33, 95% CI [-0.57, -0.08], p = 0.0097; Figure 2d), whereas selectivity showed a non-significant trend toward reduction (β = -0.04, 95% CI [-0.08, 0.002], p = 0.064; Figure 2e).

We next examined the stability of spatial coding during correct trials. At the population level, spatial stability was quantified using population-vector (PV) correlations between odd and even trials across all detected CA1 neurons within each imaging session (Figure 2f). Young 5xFAD mice showed significantly reduced PV correlations compared with controls (r_rb_ = -0.45, p = 0.031), indicating impaired population-level spatial consistency despite preserved task performance. At the single cell level, within-session tuning curve stability was also significantly lower in 5xFAD mice (β = -0.02, 95% CI [-0.04, -0.002], p = 0.029; Figure 2g), indicating reduced reproducibility of place cell spatial tuning across trajectories within the same session.

We next asked whether this instability was uniform across the session or changed as trials accumulated within session. Trial-to-trial tuning curve correlation matrices showed weaker correlations in young 5xFAD mice than in controls, particularly during the initial trials and after additional trials accumulated (Figure 2h). To quantify this pattern, we analyzed trial-to-trial curve correlations using a sliding trial window in each session. In control mice, tuning correlations were relatively stable across trials. In contrast, 5xFAD mice showed lower correlations during the initial trials, followed by a transient period in which correlations were comparable to controls, before declining again later in the session (Figure 2i, left). Analysis across trial windows supported this pattern, with reduced tuning curve correlations in 5xFAD mice during trials 1-5 (rrb = -0.47, p = 0.026) and after trial 15 (rrb = -0.28, p = 0.028), but no significant difference during trials 5-15 (rrb = -0.25, p = 0.26; Figure 2i, right). Thus, young 5xFAD mice showed reduced CA1 tuning stability during initial task trials and were less able to maintain stable spatial tuning as trials accumulated, despite a relatively preserved intermediate trial window.

Together, these findings show that, despite preserved behavioral performance, task-related CA1 spatial representations were already impaired in young 5xFAD mice, with weaker place field activity and reduced spatial stability at both the population and single cell levels. These findings prompted us to ask whether this early task-related coding vulnerability was uniformly distributed across CA1 place cells or was more pronounced in particular place cell populations.

### 3. Longitudinally stable CA1 place cells is largely preserved across learning in young 5xFAD mice

The long-term stability of hippocampal place cell representations is thought to support the spatial memory retention ^32, 33^. To assess whether the stability of spatial representations across learning was altered in young 5xFAD mice, we tracked the same CA1 neuronal populations across three task acquisition stages: early (day 6), middle (day 8), and final (after day 10) (Figure 3a). Place cells were identified at each stage, and longitudinally stable place cells were defined as neurons that (i) exhibited a place field at all three stages and (ii) showed tuning curve (TC) correlations between adjacent stages exceeding a predefined threshold (>0.51; Supplementary Figure 5c; Methods). This approach identified neurons that maintained consistent spatial representations across learning (Figure 3b-d; Supplementary Figure 5d).

**Figure 3.**
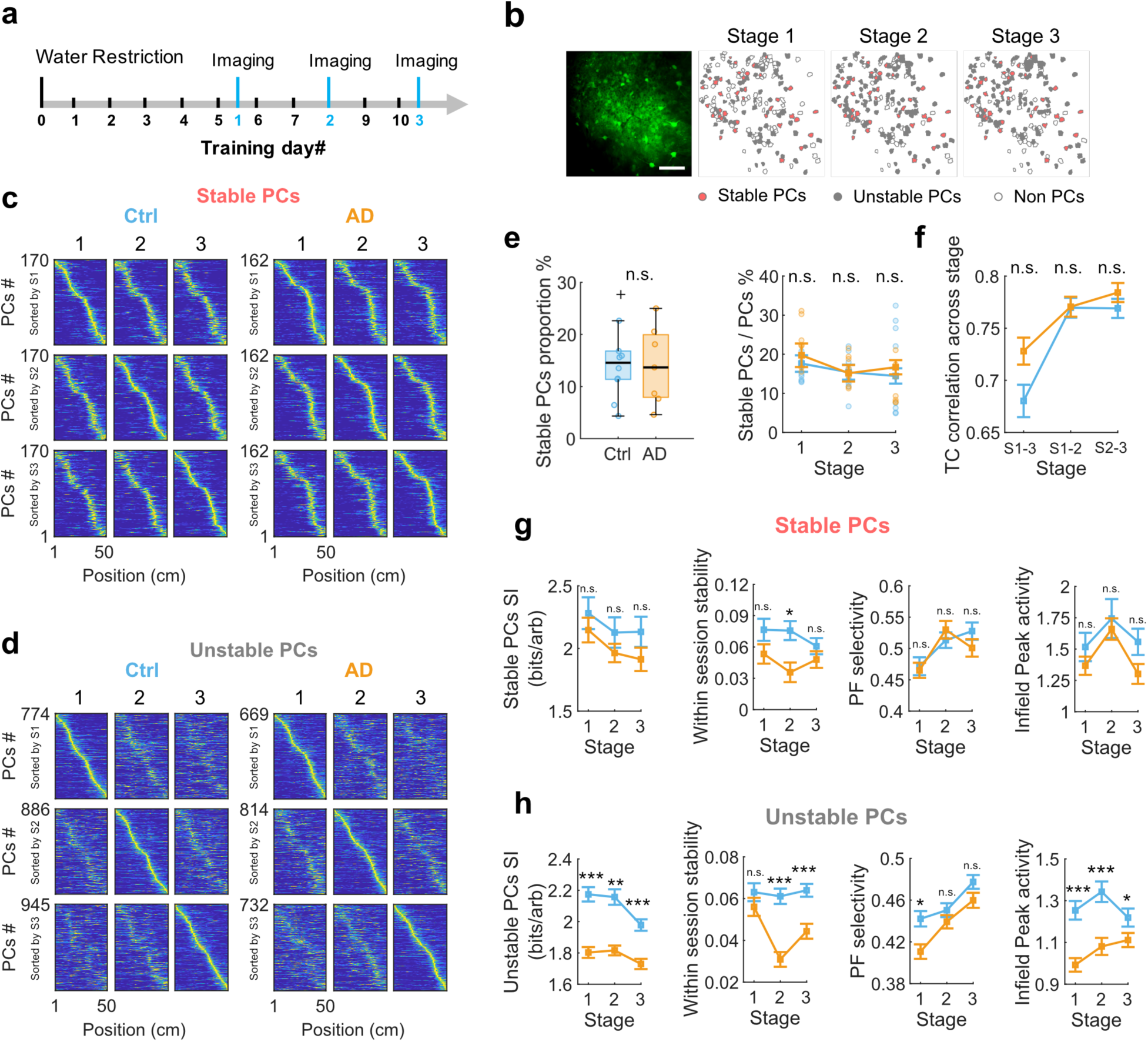
Spatial coding properties of stable and unstable CA1 place cells across learning stages in young 5xFAD mice. **a.** Experimental timeline for longitudinal imaging during task acquisition. CA1 activity was imaged at three training stages: stage 1, around training day 6; stage 2, around training day 8; and stage 3, after training day 10. **b.** Example CA1 field of view from a 5xFAD mouse, with cell classifications across the three imaging stages. Red contours indicate stable place cells, gray contours indicate unstable place cells and open contours indicate non-place cells. Scale bar, 100 µm. **c.** Linearised spatial activity maps of stable place cells across the three imaging stages in control and 5xFAD mice. Cells are sorted by peak position at each stage. **d.** Linearised spatial activity maps of unstable place cells across the three imaging stages in control and 5xFAD mice. Cells are sorted by peak position at each stage. **e.** Left, proportion of stable place cells relative to all recorded cells. Right, proportion of stable place cells relative to all place cells identified at each stage. **f.** Tuning curve correlations of stable place cells between imaging stages, shown for stage 1 versus stage 3, stage 1 versus stage 2 and stage 2 versus stage 3. **g.** Spatial coding properties of stable place cells across imaging stages, including spatial information, within session stability, place field selectivity and in-field peak activity. **h.** Same as in g, but for unstable place cells. Data are shown as box plots with overlaid session level points or as mean ± s.e.m., as indicated. Control group: 170 stable place cells and 774, 886 and 945 unstable place cells at stages 1, 2 and 3, respectively, from 10 imaging sessions in 5 mice. 5xFAD group: 162 stable place cells and 669, 814 and 732 unstable place cells at stages 1, 2 and 3, respectively, from 7 imaging sessions in 4 mice. Statistical comparisons were performed using two-sided Wilcoxon rank-sum tests with Holm-Bonferroni correction for multiple comparisons. n.s., not significant; *P < 0.05; **P < 0.01; ***P < 0.001.

Behavioral alternation performance did not differ between groups across stages (Genotype × Stage: p = 0.61, LMM; Supplementary Figure 5). The proportion of stable place cells relative to all recorded cells was similar between groups (Ctrl: 14.7 ± 2.2%; 5xFAD: 14.1 ± 2.9%; p = 1; Figure 3e). Likewise, the proportion of stable cells among all place cells did not differ at any stage (stage 1: rrb = 0.09, p = 0.89; stage 2: rrb = 0.06, p = 0.89; stage 3: rrb = 0.37, p = 0.23; Figure 3e). TC correlations of stable place cells between stages were unchanged (stage 1-3: r_rb_ = 0.12, p = 0.56; stage 1-2: r_rb_= 0.007, p = 0.91; stage 2-3: r_rb_ = 0.07, p = 0.25; Figure 3f).

Most within-session coding properties of longitudinally stable place cells were preserved in 5xFAD mice, including spatial information (stage 1: r_rb_ = -0.001, p = 1; stage 2: r_rb_ = - 0.07, p = 1; stage 3: r_rb_ = -0.02, p = 1; Figure 3g), in-field selectivity (stage 1: r_rb_ = -0.02, p = 1; stage 2: r_rb_ = 0.05, p = 0.70; stage 3: r_rb_ = -0.09, p = 0.68) and peak in-field activity (stage 1: r_rb_ = -0.13, p = 0.46; stage 2: r_rb_ = -0.07, p = 1; stage 3: r_rb_ = -0.12, p = 0.59). Within-session TC stability was comparable at stages 1 and 3, with a modest reduction in 5xFAD mice at stage 2 (stage 1: r_rb_ = -0.11, p = 0.72; stage 2: r_rb_ = -0.21, p = 0.02; stage 3: r_rb_ = -0.07, p = 1). Thus, the spatial coding properties of place cells with stable representations across learning were largely preserved in young 5xFAD mice.

By contrast, place cells with unstable spatial represnetations across stages showed reduced spatial information in 5xFAD mice relative to controls (stage 1: r_rb_ = -0.20, p < 0.001; stage 2: r_rb_ = -0.13, p = 0.003; stage 3: r_rb_ = -0.18, p < 0.001; Figure 3h) and lower within-session TC stability (stage 1: r_rb_ = -0.06, p = 0.16; stage 2: r_rb_ = -0.18, p < 0.001; stage 3: r_rb_ = -0.13, p < 0.001). These cells also showed reduced in-field selectivity at stage 1 (stage 1: r_rb_ = -0.10, p = 0.04; stage 2: r_rb_ = -0.04, p = 0.89; stage 3: r_rb_ = -0.07, p = 0.23) and reduced peak in-field activity across stages (stage 1: rrb = -0.22, p < 0.001; stage 2: rrb = -0.17, p < 0.001; stage 3: rrb = -0.09, p = 0.03). Together, these findings indicate that the task-related spatial coding impairment identified in young 5xFAD mice was not uniformly distributed across CA1 place cells. Coding properties were largely preserved in longitudinally stable place cells, whereas deficits were more pronounced among place cells with unstable representations across learning.

### 4. Trajectory-dependent contextual coding is weakened in young 5xFAD mice

Having identified greater coding impairments among place cells with longitudinally unstable representations across learning, we next asked whether a more task-specific dimension of CA1 representation, trajectory-dependent contextual coding, was also altered in young 5xFAD mice. We focused on splitter cells, which show differential activity on the common stem depending on the previous or upcoming trajectory and are thought to encode task-relevant contextual information and contribute to flexible memory-guided behavior ^34, 35^. To identify splitter cells, we examined neurons with a place field spanning at least five spatial bins along the stem arm and exhibiting trajectory-dependent activity during left and right-turn trials (Figure 4a; Methods). For each neuron, spatial tuning curves were constructed separately for correct left-right (LR) and right-left (RL) trials. Trajectories were classified as high-rate or low-rate based on which turn direction produced the higher peak spatial activity on the stem arm (Figure 4b). Splitter cells were identified based on significant differences between high- and low-rate tuning curves, and trajectory-dependent coding was further quantified using several complementary metrics (Methods).

**Figure 4.**
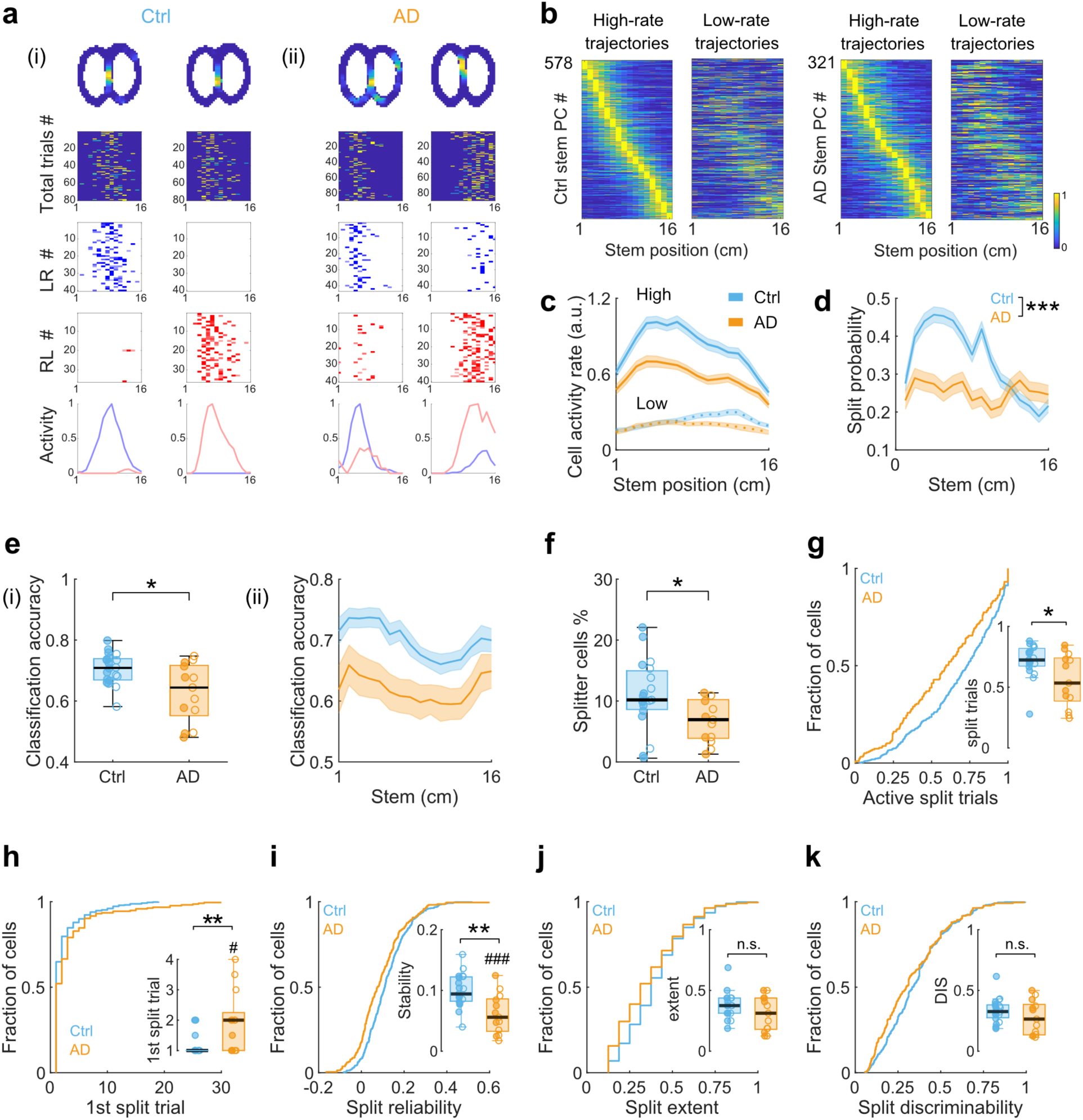
Weakened trajectory-dependent contextual coding in young 5xFAD mice. **a.** Representative stem place cells exhibiting trajectory-dependent splitting activity in control and 5xFAD mice. For each example, panels show, from top to bottom, normalized two-dimensional calcium activity maps, calcium activity across all stem trials, calcium activity during left-to-right trials, calcium activity during right-to-left trials, and normalized tuning curves for the two turn directions. **b.** Spatial activity maps of stem place cells on high-rate and low-rate trajectories. For each cell, LR and RL trajectories were classified as high-rate or low-rate according to the trajectory with the higher peak activity on the stem. Activity was normalized to each cell’s peak activity and cells were sorted by field position on the high-rate trajectory. **c.** Mean spatial activity along the common stem for high-rate and low-rate trajectories. Shaded areas indicate mean ± s.e.m. **d.** Probability of individual stem bin exhibiting significant splitting activity. **e.** Trajectory-decoding accuracy of a linear discriminant analysis classifier using all stem-active neurons. i, median decoding accuracy for each imaging session. ii, decoding accuracy along the common stem. **f.** Proportion of splitter cells among all recorded cells. One-sided Wilcoxon rank-sum test. **g-k.** Splitting properties of splitter cells. Cumulative plots show cell level distributions, and inset box plots show session level medians. **g.** Active split trial ratio. **h.** First trial exhibiting splitting activity. **i.** Trial-to-trial spatial correlation between consecutive trials on the high-rate trajectory. **j.** Split extent, quantified as the proportion of the stem place field exhibiting splitting activity. **k.** Split discriminability between high- and low-rate trajectories. Statistics in g-k were assessed using LMM test. Hash symbols indicate a genotype × sex interaction effect. Control: stem place cells, n = 578; splitter cells, n = 478; 19 imaging sessions from 9 mice. 5xFAD: stem place cells, n = 321; splitter cells, n = 217; 13 imaging sessions from 7 mice. n.s., not significant; *P < 0.05; **P < 0.01; ***P < 0.001.

Spatial tuning activity along the high-rate trajectory was reduced in young 5xFAD mice (F(1, 14352) = 214.77, p < 0.001; Figure 4c). The probability that individual stem bin exhibited significant splitting activity was also reduced in AD mice (Genotype × Spatial bin interaction: F(15, 14352) = 8.51, p < 0.001; Figure 4d). Consistently, At the population level, a linear discriminant analysis decoder trained on all stem active neurons showed significantly lower trajectory decoding accuracy in 5xFAD mice (r_rb_ = -0.49, p = 0.021; Figure 4e), indicating reduced population level trajectory information. The proportion of splitter cells was significantly lower in 5xFAD mice than in controls (Ctrl: 10.9 ± 1.3%; AD: 8.6 ± 2.4%; p = 0.037; Figure 4f).

Next, we evaluated several metrics of splitting quality. The active split-trial ratio was significantly reduced in 5xFAD mice (β = -0.12, 95% CI [-0.22, -0.02], p = 0.02; Figure 4g), and splitting activity emerged later across trials (first split trial: β = 0.98, 95% CI [0.40, 1.58], p = 0.001; Figure 4h). Trial-to-trial reliability on high-rate trajectories was also reduced (β = -0.03, 95% CI [-0.05, -0.01], p = 0.001; Figure 4i). Significant Genotype × Sex interactions were detected for first split trial and trial-to-trial reliability. By contrast, neither split extent (β = -0.04, 95% CI [-0.11, 0.016], p = 0.15; Figure 4j) nor split discriminability (β = -0.03, 95% CI [-0.09, 0.03], p = 0.34; Figure 4k) differed significantly between 5xFAD mice and controls. Together, these findings indicate that trajectory-dependent activity occurred on fewer trials, emerged later, and was less spatially reliable across trials in young 5xFAD mice, despite relatively preserved splitting strength when present.

Splitter cell quantity and coding quality were also positively associated with behavioral performance. Alternation accuracy correlated with splitter cell proportion (r = 0.48, p = 0.01), split extent (r = 0.60, p = 0.001), and split discriminability (r = 0.57, p = 0.002; Supplementary Figure 6).

To determine whether these effects reflected a general impairment of spatial coding on the common stem, we separately analyzed non-splitter stem place cells. Neither the proportion nor the trial-to-trial spatial reliability of non-splitter stem place cells was significantly reduced in 5xFAD mice (Supplementary Figure 7). Moreover, the overall proportion of stem place cells was also not associated with behavioral performance. Together, these findings indicate a selective impairment of trajectory-dependent contextual coding, rather than a general deficit in spatial coding on the common stem.

### 5. Trajectory-dependent contextual encoding patterns is reorganised in young 5xFAD mice

Having established that trajectory-dependent contextual representations were weakened in 5xFAD mice, we next asked whether the type of trajectory information represented by the remaining splitter cells was also altered. Specifically, we examined whether splitter cell activity preferentially reflected past or upcoming trajectory information. We applied an dissimilarity analysis adapted from Varga et al.^36^. Error trials were required to dissociate retrospective and prospective trajectory information coding ^37^. Analyses were therefore restricted to mice with at least three left error and right error trials. For each spatial bin, we calculated dissimilarity of cell activity between relevant trial types to quantify retrospective and prospective components (e.g., LR vs. LL for retrospective; LR vs. RR for prospective; Figure 5a,b; Methods). Lower dissimilarity indicated greater similarity between trials sharing the corresponding past or upcoming trajectory. These measures were then used to derive a Retro-Pro bias index for each cell (Figure 5c). Consistent with previous studies ^36, 37^, most splitter cells in our task exhibited a strong retrospective bias (Figure 5d).

**Figure 5.**
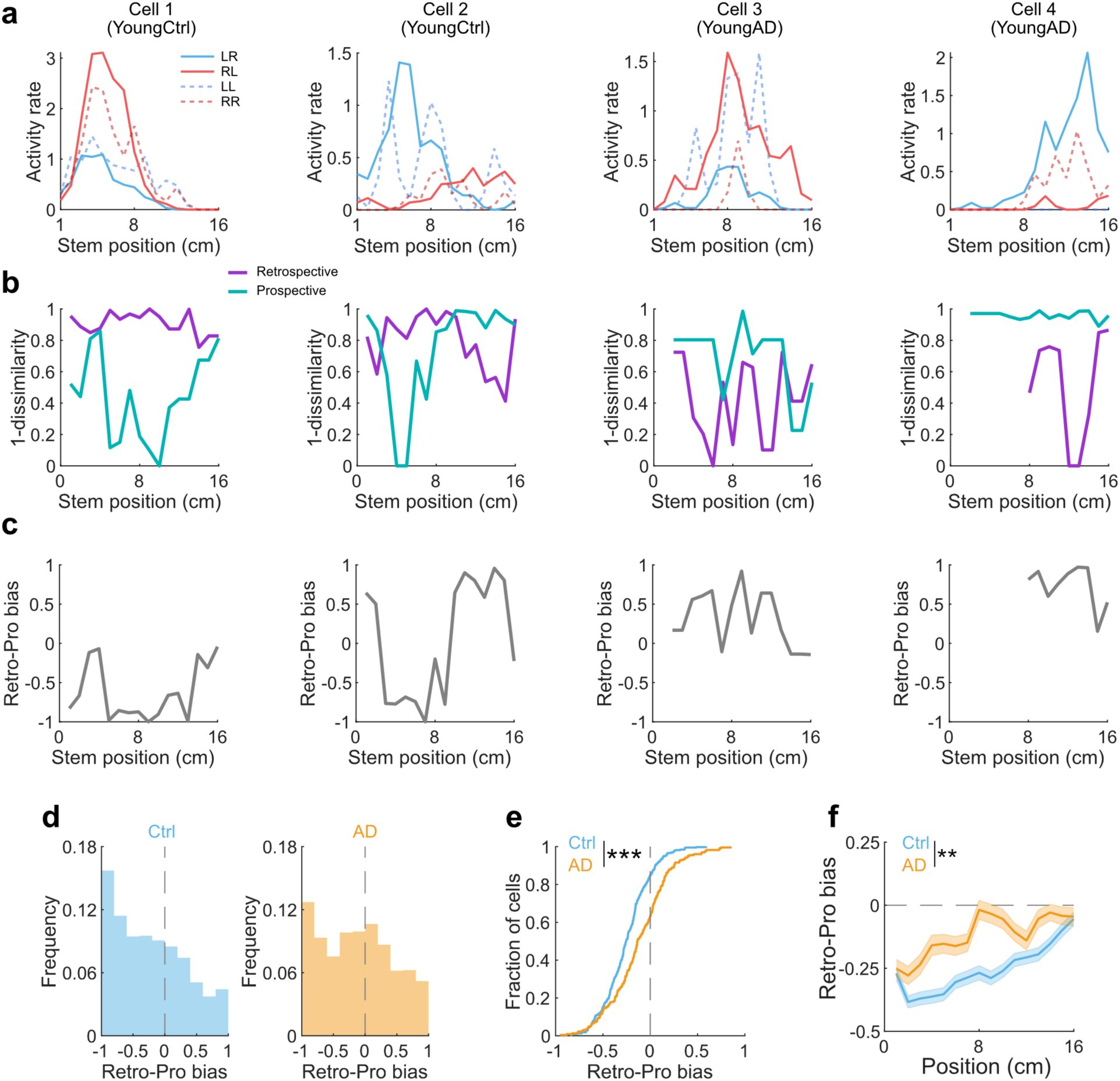
Altered hippocampal trajectory-dependent encoding patterns in young 5xFAD mice. **a.** Example splitter cells from young control and young 5xFAD mice. Spatial activity rate curves are shown along common stem for left-to-right (LR), right-to-left (RL), left-to-left (LL) and right-to-right (RR) trial types. **b.** Retrospective and prospective trajectory similarity for the example cells shown in panel **a.** Similarity was calculated as 1 − normalized dissimilarity, where dissimilarity was defined as −log10(P) from a Wilcoxon rank-sum test comparing cell activity between relevant trial types at each spatial bin (e.g., LR versus LL and RL versus RR for retrospective coding and LR versus RR and RL versus LL for prospective coding). **c.** Retro-Pro bias index along stem arm for the example cells shown in panel **a**, ranging from −1 (purely retrospective encoding) to 1 (purely prospective encoding). **d.** Histogram distribution of Retro-Pro bias values in control and AD mice. **e.** Cumulative distribution of mean Retro-Pro bias across splitter cells. The dashed line indicates zero bias. **f.** Retro-Pro bias along the common stem, from 16-0 cm toward the choice point. Shaded regions indicate mean ± s.e.m. across cells. Ctrl: 384 splitter cells from 10 imaging sessions in 5 mice. 5xFAD: 225 splitter cells from 7 imaging sessions in 4 mice. Two-way ANOVA for the Genotype × Spatial Bin interaction. **P < 0.01; ***P < 0.001.

Despite preserved alternation performance at this age, young 5xFAD mice showed marked shift in contextual encoding types. The Retro-Pro bias was significantly shifted toward more prospective-biased encoding compared with controls (β = 0.13, 95% CI [0.09, 0.18], p < 0.001; Figure 5e). This shift was characterized by a reduced retrospective component and an increased prospective component in 5xFAD mice (Supplementary Figure 8). Furthermore, whereas control mice gradually transitioned toward prospective encoding only when approaching the choice point, 5xFAD mice showed an earlier shift toward to prospective encoding along the stem (Genotype × spatial bin interaction: F(15, 8779) = 2.29, p = 0.003; two-way ANOVA; Figure 5f). Together, these findings indicate that the balance between past- and future-related trajectory information is reorganised in young 5xFAD mice.

### 6. Medial spetal cholinergic activation improved CA1 coding in young 5xFAD mice

Given the established role of medial septal cholinergic input in modulating hippocampal function and spatial coding^38^, we next asked whether the early CA1 coding abnormalities identified in young 5xFAD mice remained acutely modulable by cholinergic activation. To selectively activate medial spetal cholinergic neurons, AAV-hSyn-DIO-hM3D(Gq)-mCherry was injected into the medial septum of ChAT-Cre^+^/5xFAD^+^ (AD) and ChAT-Cre^+^/5xFAD^-^ (control) mice, and AAV-hsyn-jGCaMP7s was injected in dorsal CA1. A cranial window was then implanted for in vivo calcium imaging (Figure 6a). After all mice reached the task acquisition criterion, vehicle (saline) was administered intraperitoneally on the first imaging day and CNO on the second imaging day. Thus, chemogenetic activation was performed after task acquisition and did not test effects on task learning. CNO administration also did not alter behavioral alternation performance or locomotion in either control or 5xFAD mice (Ctrl: p = 0.81; 5xFAD: p = 0.64; two-sided Wilcoxon signed-rank test; Figure 6b and Supplementary Figure 9a,b).

**Figure 6.**
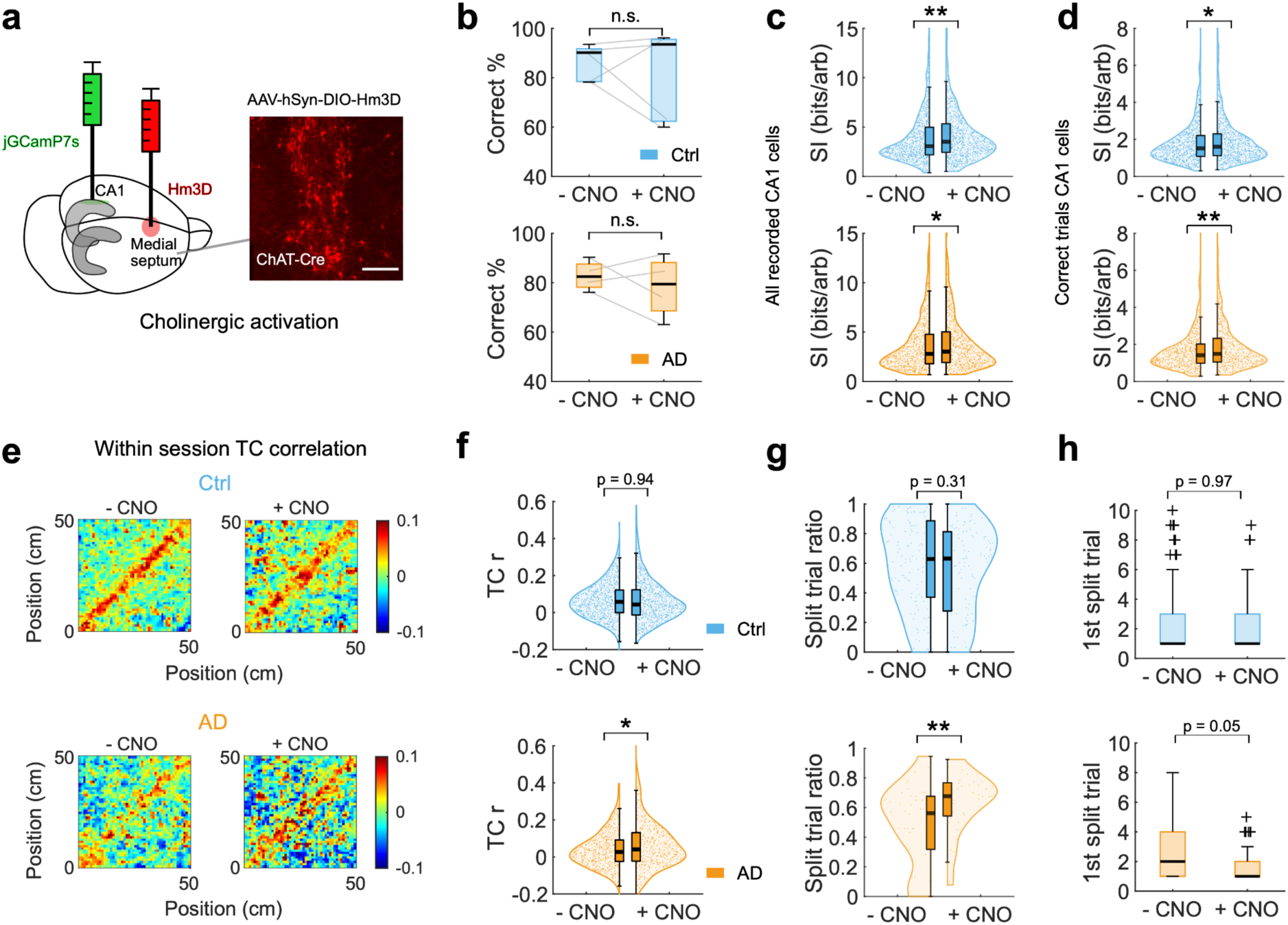
Medial septal cholinergic activation modulates CA1 spatial and contextual coding in young 5xFAD mice. **a.** Experimental schematic showing DREADD (hM3Dq) expression in medial septal cholinergic neurons and jGCaMP7s expression in hippocampal CA1. Right: representative image of hM3Dq expression in the medial septum ChAT-Cre mouse. Scale bar, 100 µm. **b.** Alternation performance following intraperitoneal (i.p.) administration of saline (-CNO) and clozapine-N-oxide (+CNO) in control and 5xFAD mice. Two-sided Wilcoxon signed-rank tests. **c.** Global spatial information of all tracked CA1 neurons after vehicle and CNO administration. **d.** Spatial information of CA1 place cells during correct trials following vehicle and CNO treatment. **e-f.** WIthinin-session stability of CA1 place cells during correct trials under vehicle and CNO administration. Mean within-session stability of CA1 place cells. **g.** Active split-trial ratio of stem place cells. **h.** First trial exhibiting trajectory-dependent splitting activity in stem place cells. For clarity, outliers >10 are not shown. Data in panels **c, d, f-h** were analyzed using linear mixed-effects models with treatment (CNO versus vehicle) as a fixed effect and mouse identity as a random intercept. Ctrl, 964 detected neurons from 5 mice; 697 and 522 place cells, including 140 and 122 stem place cells, following vehicle and CNO administration, respectively. 5xFAD, 777 detected neurons from 4 mice, 522 and 449 place cells, including 46 and 41 stem place cells, following vehicle and CNO administration, respectively. n.s., not significant; *P < 0.05; **P < 0.01.

Tracking the same neuronal populations across vehicle and CNO sessions, we found that global spatial information, computed from two-dimensional spatial firing maps, increased following CNO administration in both control (β = 0.37, 95% CI [0.13, 0.61], p = 0.002) and 5xFAD mice (β = 0.30, 95% CI [0.06, 0.54], p = 0.013). Spatial information of place cells during correct trials similarly increased in both groups (Ctrl: β = 0.12, 95% CI [0.008, 0.23], p = 0.036; AD: β = 0.19, 95% CI [0.06, 0.32], p = 0.005). In 5xFAD mice, within-session stability of place cell tuning also increased following CNO administration (β = 0.02, 95% CI [0.004, 0.036], p = 0.015; Figure 6e,f), whereas no significant change was observed in control mice (β = 0.00, 95% CI [-0.01, 0.01], p = 0.95). However, peak in-field activity remained reduced in 5xFAD mice following CNO administration, and place field selectivity was not significantly altered (Supplementary Figure 9c,d). A reduction in place-field size was observed in control mice after CNO administration (Supplementary Figure 9e).

We next assessed trajectory-dependent activity in stem place cells. The active split-trial ratio increased significantly in 5xFAD mice following CNO administration (β = 0.15, 95% CI [0.05, 0.26], p = 0.003; Figure 6g), indicating an increase in the proportion of trials showing trajectory-dependent activity. Moreover, the first-trial onset of splitting activity exhibited a trend toward earlier emergence in 5xFAD mice (β = -1.25, 95% CI [-2.50, 0], p = 0.050; Figure 6h), whereas no significant changes were observed in control mice (β = -0.02, 95% CI [-1.46, 1.40], p = 0.97). Other measures of trajectory-dependent coding, including split extent, trial-to-trial reliability, and split discriminability, were not significantly altered by CNO in 5xFAD mice (Supplementary Figure 8g-i).

Together, these findings show that chemogenetic activation of medial septal cholinergic neurons enhanced spatial information and within-session tuning stability in CA1 place cells, while increasing the trial wise occurrence of trajectory-dependent activity. These effects were selective and did not restore all altered coding properties in young 5xFAD mice.

## Discussion

In this study, we developed a head-fixed behavioral paradigm to examine hippocampal spatial and contextual representations during a continuous alternation task in a real-world maze. Combined with in vivo two-photon calcium imaging, this approach allowed us to assess CA1 population activity under the same goal-directed behavioral demands across disease stages. Previous research has shown that the 5xFAD mice begin to develop amyloid pathology between 2 to 4 months of age, with intraneuronal accumulation of Aβ42 starting as early as 1.5 months ^27^. Amyloid burden progressively increases with age^27, 39^, and by 4 to 6 months, substantial amyloid deposition is accompanied by deficits in synaptic transmission, LTP, and sharp wave-ripple activity ^24, 27, 30, 40^. Although mild cognitive abnormalities have been reported at relatively young ages, more pronounced hippocampal-depensent behavioral deficits generally emerge with disease progression ^25, 41, 42^. Consistent with these findings, young 5xFAD mice aged 2-4 months showed preserved alternation performance in our Phi-maze task during the head-fixed alternation task, whereas clear learning impairments were evident in mice older than 7 months (Figure 1).

Disruptions in hippocampal spatial representations have been reported in both human AD and across several AD mouse models ^43, 44^. Recent longitudinal calcium-imaging work further showed that CA1 spatial tuning becomes unreliable in young 5xFAD mice before object-location memory deficits emerge^18^. Our findings extend these observations by examining different levels of CA1 representation within the same ongoing goal-directed task. Global spatial information and overall place cell proportion were relatively preserved in young 5xFAD mice, whereas place cell representations examined specifically during correct trials showed reduced peak in-field activity and impaired within-session spatial stability at both population and single-cell levels (Figure 2). Trial-to-trial analysis further revealed reduced tuning correlations during the initial trials and again as trials accumulated, with a relatively preserved intermediate period. This reduction in trial-to-trial stability may reflect reduced engagement of task-related hippocampal representations in young 5xFAD mice, consistent with evidence that hippocampal place codes are strongly modulated by behavioral engagement ^45^. Somatic calcium transient decay kinetics were comparable between young control and 5xFAD mice, suggesting that altered calcium signal kinetics are unlikely to account for the early coding abnormalities (Supplementary Figure 4).

Longitudinal tracking further showed that early coding vulnerability was not uniformly distributed across the CA1 place cell population (Figure 3). Place cells that maintained stable spatial representations across learning were largely preserved in young 5xFAD mice, including their spatial information, selectivity, peak in-field activity and within-session stability. By contrast, cells with unstable representations across stages showed more pronounced impairments in these measures. Previous work has linked persistent place cell representations to neuronal excitability and the stabilisation of spatial representations during learning ^46^, and the accumulation of stable place cells during learning correlates strongly with behavioral performance in healthy mice ^47^. The stabilisation of place cell coding is closely associated with memory demands and learning in goal-directed tasks ^48, 49^. Thus, the relative preservation of this stable place cell population, whose proportion was not reduced in young 5xFAD mice, may contribute to maintaining representational continuity during repeated training in a familiar environment.

This selective vulnerability extended to trajectory-dependent contextual coding (Figure 4, Supplementary Figure7). The hippocampus supports representations that extend beyond spatial position alone ^50, 51, 52^. Splitter cells differentiate identical locations according to preceding or upcoming trajectories and are thought to contribute to flexible memory-guided behavior ^35^. In young 5xFAD mice, trajectory-dependent coding was weakened at both population and single cell levels. Trajectory-decoding accuracy and splitter cell proportion were reduced, while trajectory-dependent activity was detected on fewer trials, emerged later, and showed lower trial-to-trial spatial reliability. By contrast, split extent and discriminability were relatively preserved. Moreover, non-splitter stem place cells did not show comparable reductions in either proportion or spatial reliability, and the overall proportion of stem place cells was not associated with behavioral performance. These findings indicate a selective impairment of trajectory-dependent contextual coding rather than a general deficit in spatial coding on the common stem.

The contextual code was not only weakened but also reorganised in the balance of past-and future-related information it carried. In young 5xFAD mice, the retrospective component was reduced and the prospective component increased, resulting in a shift toward more prospective coding (Figure 5, Supplementary Figure8). This shift also occurred earlier along the common stem in 5xFAD, whereas control mice gradually shifted toward prospective coding as they approached the choice point. One possible interpretation is that previous trajectory information is less effectively maintained as animals progress toward the next choice, potentially linked to reduced hippocampal-prefrontal synchrony reported in 5xFAD mice ^53^. However, retrospective coding may not exclusively reflect memory for the preceding choice. The stronger retrospective bias near the choice point may also partly reflect the shorter stem used in our task compared with earlier studies ^36, 37^.

The deficits observed across spatial and contextual representations may reflect vulnerability of neural processes required for the consistent formation and maintenance of task-related representations. 5xFAD mice exhibit impaired synaptic transmission and reduced LTP from relatively early disease stages ^17, 24, 40, 54^. Changes in CA1 representations across learning rely heavily on synaptic plasticity driven by inputs from entorhinal cortex layer 3 projections ^55^. Rapid synaptic plasticity can also contribute to the emergence of splitter-cell activity ^23^. These observations provide a plausible context for the spatial and contextual coding abnormalities observed here. However, our experiments did not directly measure synaptic efficacy or plasticity, and the present findings therefore provide insight at the level of neural coding and representation rather than identifying a specific synaptic mechanism.

Basal forebrain cholinergic neurons send dense projections to the hippocampus and play a central role in regulating hippocampal synaptic plasticity, learning and memory^26^. Cholinergic activity also promotes hippocampal theta oscillations, which support spatial coding and memory-guided behavior ^56^. Medial septal cholinergic activation enhanced CA1 spatial and contextual representations in young 5xFAD mice (Figure 6). These effects were selective rather than global, as cholinergic activation improved spatial information and within-session stability without restoring the reduced peak in-field activity. Similarly, trajectory-dependent activity occurred on a greater proportion of trials, whereas split extent and discriminability remained unchanged, indicating partial modulation rather than complete restoration of early CA1 coding deficits.

Cholinergic system is known to be compromised in AD, and cholinergic decline has been linked to cognitive deterioration in several neurodegenerative conditions^57, 58, 59^. Previous work in 5xFAD mice showed that cholinergic fibre pathology precedes overt cholinergic neuronal loss and progressively involves the hippocampus, whereas basal forebrain cholinergic neuron loss becomes detectable at approximately 9 months of age^60^. However, hippocampal cholinergic innervation was not directly assessed in our cohort, and we therefore cannot determine whether the observed CA1 coding abnormalities or their modulation are related to the extent of cholinergic degeneration. Theta abnormalities have been reported in 5xFAD mice from early disease stages, and direct activation of septo-hippocampal cholinergic neurons can enhance hippocampal theta oscillations^61, 62^. Theta modulation therefore represents one possible mechanism contributing to the improvement in CA1 coding observed in our study. Although Alzheimer’s disease remains incurable, several approved treatments, including donepezil, galantamine and rivastigmine, act as cholinesterase inhibitors and increase acetylcholine levels. These observations suggest that targeting basal forebrain cholinergic neurons may still offer a promising strategy for early intervention in AD.

Early tetrode studies were limited by the number of neurons that could be recorded simultaneously, whereas two-photon imaging in virtual-reality environments, although scalable, does not reproduce all multimodal sensory cues available during real-world navigation ^63, 64^. More recent real-world head-fixed mazes address some of these constraints ^65^. Our study extends this framework by combining large-scale CA1 imaging with a goal-directed continuous alternation task that allows broad spatial, task-related spatial, and trajectory-dependent contextual representations to be examined within the same behavioral setting. Together, our findings reveal a progression from relatively preserved broad spatial coding to impaired task-related spatial representations, greater vulnerability among place cells with unstable representations across learning, and selective weakening and reorganisation of trajectory-dependent contextual coding. These early abnormalities precede overt decline in spatial alternation performance and remain partially amenable to cholinergic modulation.

### Limitations of the study

Several limitations should be considered. First, although sex was included in our statistical analyses and significant Genotype × Sex interactions were detected for several coding measures, the relatively small numbers of male and female mice limited detailed sex-stratified analyses. Given reported sex differences in AD pathology and cognitive decline in both humans and 5xFAD mice ^66, 67, 68, 69, 70, 71^, larger cohorts will be required to determine the robustness and biological significance of these interactions. Second, somatic calcium transient decay kinetics were comparable between young control and 5xFAD mice, reducing the likelihood that altered decay kinetics account for the principal early-stage coding deficits (Supplementary Figure 4). However, the shorter decay observed in older 5xFAD mice indicates that altered calcium dynamics may influence the interpretation of calcium derived activity measures at later disease stages. Third, our in vivo calcium imaging approach did not directly measure hippocampal oscillatory activity. We therefore could not determine whether theta modulation contributed to the effects of medial septal cholinergic activation on CA1 coding. Simultaneous electrophysiological recordings will be required to address this possibility. Fourth, The fixed vehicle before CNO treatment order may also have introduced potential order effects, although no differences were observed between the no-injection baseline and vehicle sessions (Supplementary Fig. 9f). Finally, our behavioral assessment was limited to a single task that lacked an explicit delay period, relying instead on the animal’s natural return time to the choice point. Incorporating a controlled delay (such as with a servomotor gate) could increase the task’s reliance on intrinsic hippocampal processing, as suggested by studies in freely moving animals, but was beyond the scope of the present work ^34^.

## Materials and Methods

### Experimental Animals

5xFAD mice express human APP and PSEN1 transgenes with a total of five AD-linked mutations: the Swedish (K670N/M671L), Florida (I716V), and London (V717I) mutations in APP, and the M146L and L286V mutations in PSEN1. Male hemizygous 5xFAD mice (B6SJL-Tg (APPSwFILon, PSEN1∗M146L∗L286V) 6799Vas/Mmjax, JAX stock # 034840) were crossed with female C57BL/6J wild type mice to generate 5xFAD positive and wild-type littermate controls. Male hemizygous 5xFAD mice were crossed with female homozygous Chat-IRES-Cre(△neo) mice (JAX stock # 031661 on a C57BL/6J genetic background) to maintain the 5xFAD positive (5xFAD^+^/Chat-Cre^+^ as 5xFAD group) and non-5xFAD (5xFAD^−^/Chat-Cre^+^ as control group) mice. Colonies under standard animal breeding at Imperial College London. The animal room was kept on a reverse 12:12 light-dark cycle daily.

Real-time PCR genotyping was performed by Transnetyx using APPswTg and huPSEN1Tg probes to identify 5xFAD transgene carriers. Animals were divided into two age groups: young mice (control, median age 3.6 months, n = 10; 5xFAD, median age 3.6 months, n = 7) and older mice (control, median age 8.6 months, n = 6; 5xFAD, median age 7.9 months, n = 6). Detailed animal information is provided in Supplementary Table 1. 5xFAD-negative littermates were used as controls. Both male and female mice were included in the study. All experimental procedures were conducted under the Animals (Scientific Procedures) Act 1986 and in accordance with UK Home Office and institutional guidelines, under Project Licences 7009095 and PP6988384 held by S.R. Schultz.

### Aβ staining

MOAB-2, clone 6C3 (Millipore), was used as a pan-Aβ antibody to assess the percentage area occupied by amyloid-β. Sagittal brain sections (30 μm) were obtained using a cryostat. Antigen retrieval was performed with 98% formic acid for 5 min, followed by permeabilisation for 20 min with 0.6% H₂O₂ in TBS containing 0.25% Triton X-100 (TBS-Tx). Free-floating sections were subsequently blocked for 1 h with 10% FBS in 0.1% TBS-Tx and incubated overnight at 4°C with anti-Aβ MOAB-2 (clone 6C3; 1:1000) in 2% FBS and 0.2% BSA in 0.02% TBS-Tx.

The following day, sections were washed and incubated with a biotinylated anti-mouse secondary antibody (1:200) in 2% FBS and 0.2% BSA in 0.1% TBS-Tx for 1.5 h at room temperature. Sections were then incubated with avidin-biotin complex (ABC; Vector) and subsequently developed with diaminobenzidine (DAB; Vector) to visualise staining. Sections were sequentially washed in ddH₂O and PBS, mounted onto slides, and left to dry overnight. The following day, slides were dehydrated through increasing concentrations of ethanol, cleared in xylene, and mounted in DPX (Sigma-Aldrich).

Slides were scanned using an APERIO AT2 scanner (Leica). The percentage of Aβ-positive area in different hippocampal subregions was quantified using HALO software from 2–3 sections per animal.

### Surgeries

Mice were anaesthetised with 1.5-3% isoflurane (Iso-Vet, Chanelle Pharma) and body temperature was maintained at 37°C. Analgesia was administered pre-operatively with Carprofen (5 mg/kg SC, Rimadyl, Zoetis) and Buprenorphine (0.07 mg/kg SC, Vetergesic, Ceva Animal Health Ltd). Anaesthetic depth was monitored every 10 minutes via the pedal flexion reflex. A small (0.5 mm) craniotomy was performed and 1 μl of a viral cocktail containing two adeno-associated viruses (pGP-AAV-syn-jGCaMP7s-WPRE, Addgene catalogue 104487-AAV9; pAAV-CAG-tdTomato, Addgene catalogue 59462-AAV9) in a 3:1 ratio was injected into the hippocampal CA1 region (coordinates from bregma, in mm: -1.3 and -1.5 DV, -1.8 ML, -2.0 AP). Ten minutes after viral injection, a 3-mm circular craniotomy was made, and the overlying cortex (including parietal cortex and parts of visual and hindlimb sensory cortex) was aspirated using a 27-gauge needle connected to a water pump until the corpus callosum fibres were exposed. A stainless-steel cannula (3 mm diameter, 1.5mm height) with a glass bottom was inserted and secured with histoacryl glue (B. Braun Surgical). The hippocampal imaging window was described by Dombeck et al. ^72^. A stainless-steel head plate was glued to the skull, centered over the craniotomy. The exposed skull outside the head plate aperture was covered with dental cement mixed with black powder paint. Postoperatively, mice received Carprofen (5 mg/kg/24 hrs) in oral water for three days. Behavioral training began after a 2-3 week recovery period.

### Behavioral training in head-fixed mice

#### Maze apparatus

We used a carbon fibre maze with attached visual and tactile cues, floating on the Neurotar tracker air table (Mobile HomeCage, Neurotar). Built-in magnets in maze allowed for tracking the animal’s position within the environment. The maze has a diameter of 32.5 cm, with a 16 cm-long stem arm. The corridor width for all arms was 5 cm. Visual cues were provided by attached phosphorescent tape, and tactile cues were added using Fiberglass Mesh Tape on the inner wall of the maze. The training cage tape was fully charged with daylight before the commencement of any training sessions.

#### Handling and habituation

Acclimatization to both the room and maze is essential for mice to become familiar with the training environment prior to official training sessions. Mice were handled at least two days before behavioral training. On the day before training began, mice freely explored the air-lifted T maze in the training room under red lighting for 10 minutes. During the first 5 training days, mice were allowed 5 minutes of free exploration before head fixation.

#### Spatial continuous rewarded alternation task

To motivate task engagement, mice were subjected to water restriction prior to the first day of training, while food was available ad libitum throughout the experiment. Body weight was monitored daily and typically stabilised at 85–90% of the initial body weight after approximately 5 days. If insufficient water was obtained during task training, supplementary water was provided in the home cage after the second daily session to maintain body weight above 85% of the initial body weight. Mice were trained in the dark during the learning of the alternation task. For the first trial, mice could freely choose reward arm. A 4 μl reward was delivered via a lick spout attached to the head-plate clamp and controlled by a peristaltic pump (Campden Instruments), accompanied by a beep. In subsequent trials, a reward was only delivered when mouse entered the reward arm opposite to the one visited in the previous trial. A complete trial required the mouse to pass through the previous trial reward position, return arm, stem arm, choice arm, and arrive at the next reward location. Head-fixed animals were trained for 45 minutes each session twice a day, with at least 4 hours between sessions. From the 6th training day onward, 1% sucrose water was used as the reward during training and imaging sessions, but still supplied water for supplementary fluid. Mouse position and behavioral data were recorded using the Neurotar Mobile HomeCage tracker (V3.0.0.64). The Phi-maze was cleaned with 70% ethanol before each mouse training or imaging recording.

#### In vivo two-photon Ca2+ imaging

Two-photon calcium imaging was conducted using a Scientifica resonant scanning microscope with a 16× Nikon water-immersion objective (LWD 0.8 NA). jGCamp7s and tdTomato fluorophores were excited at 940 nm using Ti:Sapphire laser (Mai Tai HP, Newport). The 50% (vol%) ultrasound gel was used as the immersion medium. Images (490 × 490 μm, 512 × 512 pixels) were acquired at 30.9 Hz frame rate using SciScan 1.3 software. Imaging was synchronized with behavior recording via a start TTL signal sent from the behavior recording software Neurotar. All mice underwent imaging afterwards 10th training day. Additionally, some mice were also imaged on 6th and 8th training days to enable imaging across the learning period. Each imaging session lasted no longer than two hours per mouse per day. At the beginning of each session, mice were head-fixed at the stem arm and restricted by two carbon fibre doors. Once the targeted field of view (FOV) was located, the doors were removed and imaging commenced as the mouse began running. To ensure sufficient data for spatial tuning analyses, a minimum of 80 total trials was required for each imaging session, though approximately 100 trials were usually recorded per session.

#### Methoxy-X04 labeling

To visualize amyloid plaques in vivo, mice received an intraperitoneal injection of Methoxy-X04 (10 mg/kg). Amyloid plaques were imaged in vivo in the head-fixed 5xFAD mice at least 24 h after injection using 740 nm laser excitation. At least 24 h after Methoxy-X04 injection, mice were deeply anesthetized and transcardially perfused with PBS followed by 4% paraformaldehyde. Brains were collected and imaged by ex vivo two-photon tomography using a TissueCyte imaging system (TissueVision, Newton, MA, USA). Methoxy-X04-labeled amyloid plaques were detected using 740 nm laser excitation.

#### Chemogenetic activation

For chemogenetic activation of basal forebrain cholinergic neurons, 0.5–1.0 μL AAV-hSyn-DIO-hM3D(Gq)-mCherry was injected into the medial septum (AP +0.8 mm, ML +0.1 mm, DV −4.2 to −4.0 mm from bregma), and jGCaMP7s was delivered into dorsal hippocampal CA1. DREADDs experiments were performed only after mice reached stable task performance. Animals were recorded across two consecutive imaging days. On the first day, mice received an intraperitoneal injection of saline (vehicle). On the second day, Clozapine-N-oxide (CNO; 1 mg/kg, i.p.; water-soluble CNO dihydrochloride, HelloBio HB6149) was administered 40 minutes before imaging. A low CNO dose was used to minimize off-target effects, and previous studies have reported that 1 mg/kg CNO does not alter hippocampal neuronal activity without hM3D injection in ChAT-Cre mice ^73^. The same neuronal population was tracked across vehicle and CNO sessions.

### Quantification and statistical analysis

#### Behavior data analysis

Only complete trials (see behavior training) were included in the analysis. As mice received a reward on their initial free choice, this first choice was excluded from the calculation of correct trials. Alternation performance was defined as the percentage of correct trials out of the total number of trials. To examine changes in performance across training days, we fitted a linear mixed-effects model (LMM): data ∼ 1 + Genotype × Day + (1 | MouseID). Performance (percentage of correct trials) was the dependent variable. The model included Genotype (5xFAD, Control), Training days (as a continuous numerical predictor), and their interaction as fixed effects. To account for repeated measures from the same subject, Mouse ID was included as a random intercept. To further assess learning dynamics independent of daily variability, trials from all sessions were concatenated and binned into consecutive 50-trial blocks, with the percentage of correct responses computed for each block. Learning trajectories across groups were then modeled using a quadratic mixed-effects model: Performance ∼ Genotype × (Block + Block²) + (1|MouseID). Differences in task learning performance between 5xFAD mice and their age-matched controls were determined based on the Genotype × Day and Genotype × Block interaction effects. Task acquisition criterion is the percentage of correct trials is over 70% over four consecutive trial blcoks.

#### Trajectory linearisation

To linearise the mouse trajectory, the maze was segmented into four functional zones: the return arm, stem arm, choice arm and reward arm. Trajectories were parsed accordingly, starting from the return arm, then moving up the stem arm into the choice arm, and ending in the reward arm. The reward spout marked the boundary between the choice and reward arms. Each trajectory was then categorised into L-R, R-L, L-L and R-R trial types based on maze segmentation. Linearised trajectories were mapped by trial type onto a standardised 49.5 cm track, corresponding to the estimated length of a complete trial. Trials were chronologically ordered based on the sequence of completion in the maze.

#### Imaging data processing

To extract calcium activity from individual neurons, we employed a customised image processing pipeline based on the CaImAn MATLAB package ^74^, following the protocol described by Go et al. ^65^. Imaging data were initially corrected for motion artefacts using sequential rigid and non-rigid registration methods, aligning frames a reference template from the same imaging session. Regions of interest (ROIs) were then identified, with excluding any overlapping ROIs. To enable tracking of ROIs across multiple imaging stages, images from different stages were registered to a template from one session, and ROIs were identified on a temporally concatenated video. For quantification of single-neuron calcium amplitude trains, neural activity was deconvolved using a sparse non-negative deconvolution approach. This was implemented with an exponential kernel corresponding to the decay timescale of the calcium indicator employed, utilizing the OASIS (Online Active Set method to Infer Spikes) algorithm to infer fluorescence activity into calcium trains with both onset times and amplitudes.

To eliminate out any discrepancies in OASIS event amplitude between different mice and FOVs, we fitted the distribution of event amplitudes for each FOV in each mouse to a lognormal distribution and standardised the distribution for each FOV to unit mean. This eliminates variability in event amplitude due to differences in signal to noise ratio (for instance due to imaging depth and viral expression), and makes it possible to compare calcium activity rates objectively across FOVs and animals. The resulting deconvolved calcium event amplitudes are expressed in consistent arbitrary units across the entire dataset.

#### Spatial information

To quantify spatial representations in CA1 neurons, spatial information was calculated from two-dimensional spatial rate maps for each detected neuron. The Phi-maze was discretised into 1 cm-wide bins, forming a 32 × 32 spatial grid. For each bin, the spatial activity rate was computed as the sum of standardised deconvolved calcium amplitudes recorded within that bin, divided by the total occupancy time, considering only periods when the mouse’s running speed exceeded 8 mm/s. The resulting spatial rate maps were smoothed using a Gaussian kernel with a sigma of [3, 3]. The spatial information content of each neuron was calculated using Skaggs’s spatial information method ^75^ (bits per arb, standardised amplitude rate), is calculated using the following formula:

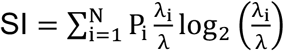

where SI is the spatial information in bits/arb. P_i_ is the probability of the mouse being in the i-th bin. λ_i_ is the mean standardised amplitude rate of the cell when the mouse in the i-th bin. λ is the overall mean standardised amplitude rate of the cell across the entire environment. N is the total number of spatial bins.

#### Population vector correlation

To assess the stability of spatial representations at populational level, we calculated population vector (PV) correlations between odd and even trials. For each session, we first constructed spatially binned activity curves for all detected cells. The population vector at each spatial bin was defined as a vector containing the calcium activity rates of all cells within that bin. We then computed the PV correlation matrix by comparing population vectors between odd and even trials across all spatial bins by Pearson correlation. The diagonal elements of this matrix represent the correlation coefficients for the same spatial positions between trial types, reflecting the stability of spatial coding along the linearized trajectory. Finally, we averaged the diagonal PV correlation coefficients across animals within each experimental group.

#### Place cell identification

Place cells from two-dimension spatial maps were identified based on the following criteria: 1) A neuron’s SI must exceed the 95th percentile of the shuffled SI values. 2) the neuron should be active within the place field for more than 2% of the time the mouse spends in that field. 3) the size of the place field must exceed 5 cm^2^. The boundary of a place field was determined by encompassing all connected occupancy bins where was at least 50% of the peak rate in the occupancy bin.

For place cells selection from one-dimension spatial maps, we calculated for correct trials (R-L and L-R) and error trials (L-L and R-R) based on same type of linearised trajactory. The criterions for 1D place cells : 1) A neuron’s SI must exceed the 95th percentile of the shuffled SI values. 2) the neuron should be active within the place field for more than 1% of the time the mouse spends in that field. 3) the neuron were active more than 30% of total number of trials.

#### Place cell quality metrics

1. In-field selectivity: the in-field and out-of-field activity rates were computed and normalized by the number of spatial bins. Selectivity was defined as (In-field – Out-field)/( In-field + Out-field). ranging from −1 (purely out-of-field firing) to 1 (purely in-field firing).
2. Peak in-field activity was defined as the maximum activity rate within each place field, corresponding to the spatial bin with the highest activity rate. For place cells with multiple place fields, in-field selectivity and peak in-field activity were calculated separately for each field and then averaged across fields to obtain one value per cell.
3. Within-session stability. To quantify the reproducibility of spatial tuning within an imaging session, the same type of correct trials were divided into odd and even subsets, with trial-wise spatial activity retained for each individual trial. For each spatial bin, activity vectors were constructed across odd and even trials, and Pearson correlations were calculated between spatial bins across the two trial subsets. The diagonal elements of the resulting correlation matrix, representing correlations between matched spatial positions, were averaged to obtain a single within-session stability value for each cell.
4. Trial-to-trial tuning correlation. To assess changes in spatial tuning across individual trials, a single trial tuning curve was constructed for each correct trial from spatially binned calcium activity along the linearised trajectory. Pearson correlations were calculated between tuning curves from individual trial pairs to generate a trial-by-trial correlation matrix for each place cell. These pairwise correlations were used to assess tuning stability as a function of trial progression and in sliding trial windows.

#### Tuning curve (TC) correlation across imaging stages

TC correlations were calculated for longitudinally tracked neurons classified as place cells at the relevant imaging stages. For each stage, calcium activity was spatially binned along the linearised trajectory and averaged across trials to generate an averaged spatial tuning curve for each place cell. Pearson correlation coefficients were then calculated between tuning curves from different imaging stages to quantify the similarity of spatial representations across learning. Correlations were calculated for stage 1 versus stage 2, stage 2 versus stage 3, and stage 1 versus stage 3.

#### Identification of longitudinally stable place cells across stages

Stable place cells were defined as neurons that (i) exhibited a place field at all three imaging stages and (ii) showed tuning-curve (TC) correlations between adjacent stages above a predefined threshold. To define this threshold, we used young control place cells as the reference population. For each place cell, odd-trial and even-trial tuning curves were first averaged across trials within the same imaging session. Real within-day TC correlations were then calculated by correlating the trial-averaged odd-trial and even-trial tuning curves from the same cell. A null distribution was generated by randomly pairing trial-averaged odd- and even-trial tuning curves from different young control place cells. This randomization was repeated 1,000 times across 500 shuffles. The stability threshold was defined as the median 95th percentile of the shuffled distribution (TC correlation = 0.51) and was applied uniformly across groups. (Supplementary Figure 5c).

#### Splitter cell identification

To identify hippocampal splitter cells, the stem arm of the figure-8 maze was divided into 1-cm spatial bins. Place cells with place fields spanning at least five consecutive bins along the stem were included for analysis. For each cell, neural activity was computed at each stem bin across trials, and spatial tuning curves were constructed separately for correct left-turn and right-turn trials. The difference between the two tuning curves was then calculated at each bin. Significance was assessed using a permutation procedure in which left- and right-turn trial labels were randomly shuffled 1,000 times. A cell was classified as a splitter cell if (i) it possessed a place field covering at least five stem-arm bins and (ii) its real tuning-curve difference exceeded the 95th percentile of the shuffled distribution in at least one spatial bin.

To ensure that splitting activity did not arise from stereotyped speed differences along the stem, we evaluated whether each spatial bin exhibited speed-independent splitting. For each neuron, we first constructed spatially binned activity for every trial. We then performed a linear mixed-effects analysis (MATLAB fitlme) for each stem place cell (defined as cells with a place field > 5 cm along the stem). The model included trial type (left/right) as a categorical predictor and animal speed as a continuous predictor, with spatially binned activity as the dependent variable. A spatial bin was considered to exhibit valid splitting activity if trial type remained a significant predictor of neuronal activity after controlling for speed. Neurons with at least two spatial bin meeting this criterion were classified as splitter cells.

#### Splitter cell quality metrics

The quality of neuronal splitting activity was assessed using five metrics, computed for identified splitter cells:

1) Split extent ^76^

The proportion of spatial bins along the stem arm that exhibited significant splitting activity.

2) Split discriminability.

Split discriminability was calculated as the sum of the absolute differences between left-and right-turn tuning curves across all stem bins, normalized by the total summed activity of both tuning curves across those bins.

3) Split reliability (Trial-to-trial correlation)

This metric quantified the consistency of splitting activity across trials. For each cell, Pearson correlation coefficients were computed between the spatial activity profiles of adjacent trials in the neuron’s preferred trajectory direction (e.g., between consecutive left–right trials if the cell preferred the L-R trajectories). Correlations were calculated only within the same imaging session.

4) Active split-trial ratio.

The proportion of trials in which a splitter cell was active during its high-rate trajectory direction.

5) First split trial

The trial number at which a cell first is active on its splitting spatial bin.

#### Discriminant classification

A linear discriminant classifier was trained using 70% of randomly selected correct trials from each imaging session. The classifier was implemented using MATLAB’s built-in function fitcdiscr. For each neuron, calcium activity at each time point while the mouse occupied the stem arm served as input features, and the mouse’s upcoming turn direction was used as the response variable. Only correct trials (left-right and right-left) were included in both the training and testing sets. The trained classifier was then used to predict turn direction on the remaining 30% of correct trials. This procedure was repeated 1,000 times, each with different randomly selected training and testing subsets. Decoding accuracy was computed within 1 cm spatial bins along the stem arm, and session level decoding accuracy was defined as the mean accuracy averaged across all spatial bins.

#### Quantification of Retrospective-Prospective Bias

We quantified the influence of past versus future behavioral choices on neuronal activity by computing a dissimilarity-based Retrospective-Prospective (RP) bias index, adapted from previously established approaches Varga et al.^36^. Analysis was restricted to recording sessions containing at least three error left trials and three error right trials. For each spatial bin along the stem, we extracted calcium activitity rates from four trial types: RL: correct left; LR: correct right; LL: error left; RR: error right. For each spatial bin, dissimilarity between two trial types was computed as: D = −log10(p), where p is the Wilcoxon rank-sum test p-value comparing firing-rate distributions across trials. Higher dissimilarity values indicate greater difference in neural activity patterns between the two compared trial types. Four pairwise comparisons were performed: Retrospective comparisons (LL vs. LR; RR vs. RL), Prospective comparisons (LL vs. RL; RR vs. LR). For each spatial bin, we derived: D_retro_=min[D(LL,LR),D(RR,RL)]; D_pro_ =min[D(LL,RL),D(RR,LR)]. The Retro–Pro (RP) bias index was then computed as: (D_retro_ - D_pro_)/ (D_retro_ + D_pro_). This value ranges from −1 (purely retrospective encoding) to +1 (purely prospective encoding), with values near zero indicating balanced encoding.

#### Statistical tests

All statistical analyses and visualisation were performed in MATLAB. Data are presented either as mean ± s.e.m. or as box plots. The significance threshold was set at p ≤ 0.05. Behavioral performance across training days or trial blocks was analyzed using linear mixed-effects models (LMMs). Genotype, Day (or Block) and their interaction (Genotype × Day/Block) were treated as fixed effects, and Mouse ID was included as a random intercept to account for repeated measures. For datasets with multiple imaging sessions per mouse (where each session corresponded to a distinct imaging field of view, with no repeated cells across sessions), linear mixed-effects models included nested random intercepts for variability (data ∼ 1 + Genotype + (1 | MouseID:SessionID)). For datasets in which each mouse contributed only a single imaging session (e.g., DREADD experiments), only MouseID was included as a random effect, as SessionID and MouseID were equivalent in this context. Stable place cells across learning stages were pooled across mice and analyzed using non-parametric comparisons. For non-parametric comparisons, effect sizes were reported as rank-biserial correlation (r_rb_). For linear mixed-effects models, estimates are reported with 95% confidence intervals. For all other comparisons, non-parametric Wilcoxon rank-sum or Wilcoxon signed-rank tests were used to avoid assumptions of normality, unless otherwise specified in the relevant methods. Data points are shown whenever possible, and Bonferroni corrections were applied for multiple comparisons. For datasets displayed as box plots, boxes indicate the interquartile range (IQR; 25th–75th percentiles), the central line denotes the median, and whiskers extend to 1.5 × IQR; points beyond this range are plotted as outliers.

## Acknowledgments

We thank Yu Liu and Nawal Zabouri for their help with animal genotyping and daily support, Jina Song for assisting with the behavioral training protocol. We also thank Marc Busche and Julija Krupic for their helpful comments and suggestions on the manuscript.

## Funding

Mrs Anne Uren and the Michael Uren Foundation to SRS

EPSRC EP/W024020/1 and EP/W035057/1 to SRS

BBSRC BB/Y514202/1 to SRS

CZI CP-2-1-Schultz to SRS

## Author contributions

Y.L. and S.R.S. conceived and designed the study. Y.L.mainly conducted experiments and collected data under guidance and supervision of S.R.S. and M.A.G.. R.A. and M.S. performed histochemistry. J.K. and S.P. performed TissueCyte imaging and reconstruction. Y.L. H.Z. and M.G. analyzed data. Y.L., M.A.G. and H.Z. contributed to imaging data processing. Y.L. and S.R.S. wrote the paper. All authors discussed and commented on the manuscript.

## Data and materials availability

Code and data used to generate the figures described in this paper will be made available via DataDryad at the time of publication, with a DOI to be included here.

---

**Supplementary Figure 1.**
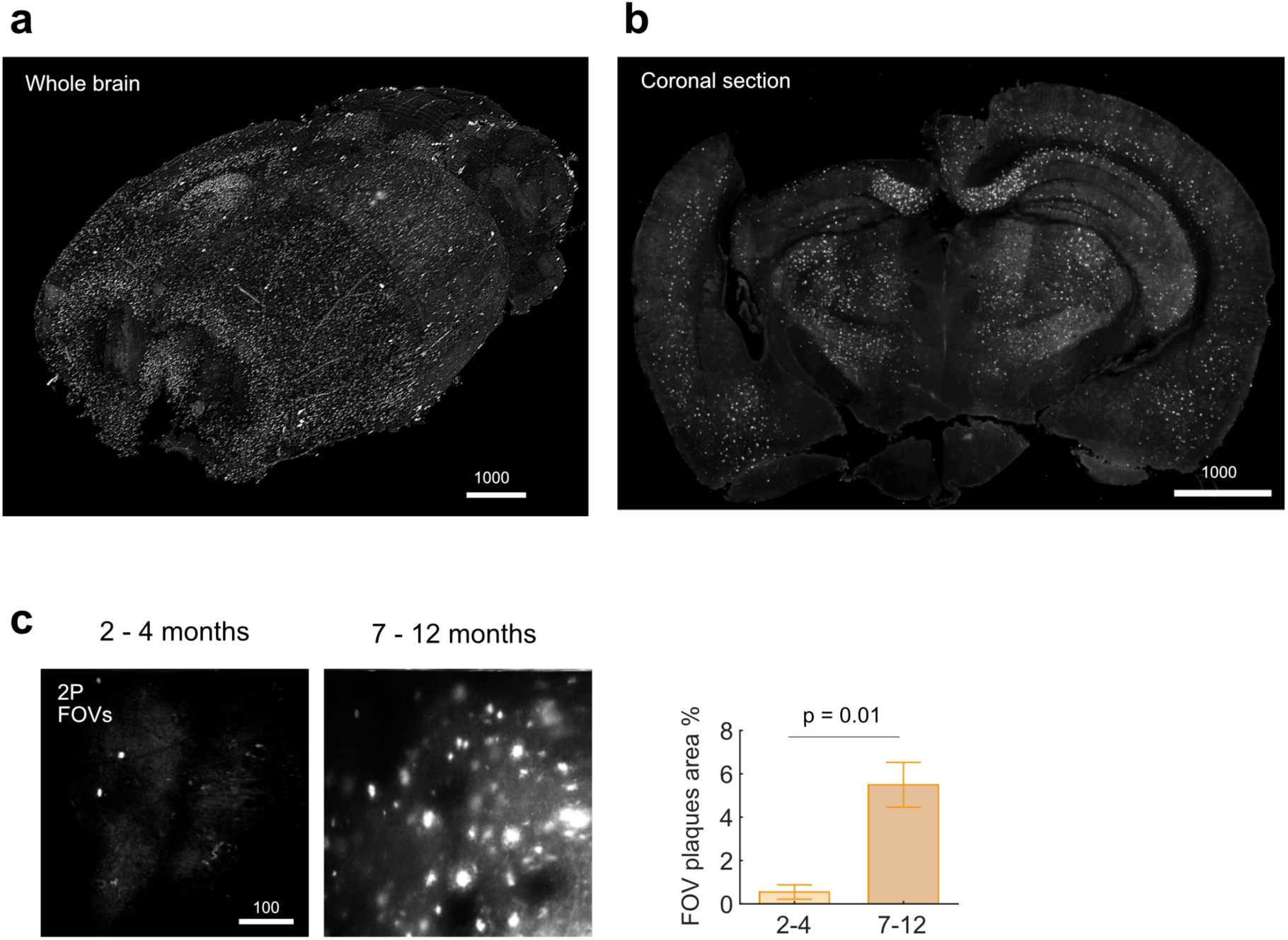
Methoxy-X04 labeled Aβ plaques in 5xFAD mice. **a.** Three-dimensional reconstruction of a 5xFAD mouse brain labeled with Methoxy-X04, from ex vivo two-photon tomography carried out with a Tissuecyte (TissueVision, Newton MA) imaging system, complementing the MOAB-2 histochemistry shown in the main text. Scale bar, 1000 µm. **b.** Representative coronal section showing Aβ deposition in the hippocampus. The tissue opening above the dorsal hippocampus corresponds to the chronic imaging window. Scale bar, 1000 µm. **c.** Representative in vivo two-photon imaging of Methoxy-X04 labeled amyloid plaques within CA1 imaging fields of view. Quantification shows the percentage of plaque-positive area relative to the total FOV area in 5xFAD mice at median ages of 3 months (n = 3 mice) and 8 months (n = 3 mice). Data are shown as mean ± s.e.m.; comparisons were performed using a two-sided Student’s t-test. Scale bar, 100 µm.

**Supplementary Figure 2.**
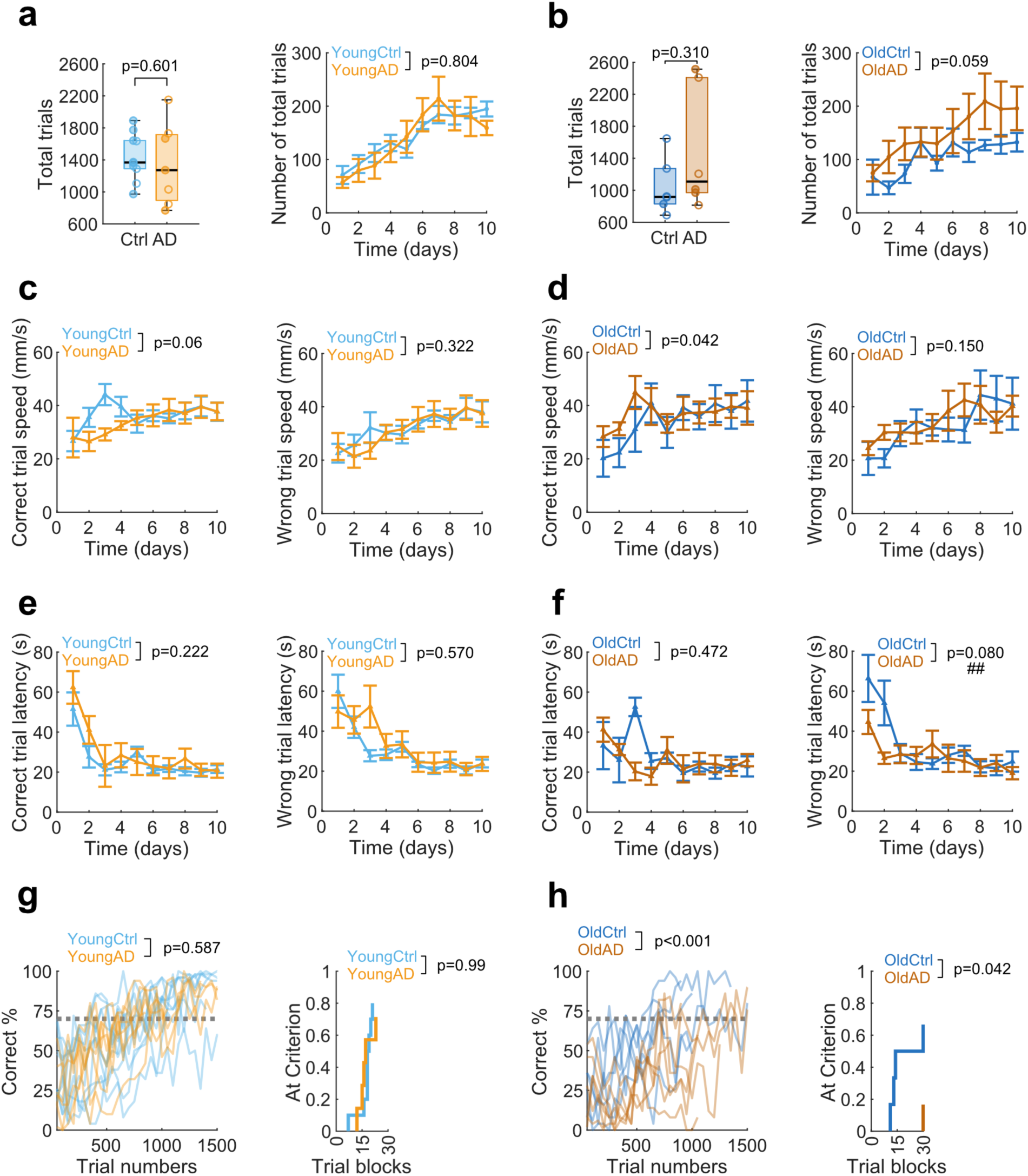
Behavioral training measures in head-fixed 5xFAD mice. **a.** Training engagement in young control and 5xFAD mice. Left, total number of trials completed across training. Right, mean number of trials completed per training day. **b.** Same as in panel a, but for old control and 5xFAD mice. **c.** Running speed across training days in young mice, shown separately for correct trials (left) and error trials (right). **d.** Same as in panel c, but for old mice. **e.** Trial latency across training days in young mice, shown separately for correct trials (left) and error trials (right). **f.** Same as in panel e, but for old mice. **g.** Trial-block analysis of alternation performance in young mice. Left, individual mouse performance across consecutive 50-trial blocks. The dashed line indicates the 70% acquisition criterion. Right, cumulative proportion of mice reaching criterion across trial blocks. **h.** Same as in panel g, but for old mice. Data are shown as mean ± s.e.m. unless otherwise indicated. P values indicate genotype effects. Hash symbols indicate significant genotype × sex interactions. Statistical comparisons were performed using linear mixed effects models for training day analyses, two-sided Wilcoxon rank-sum tests for total trial counts, and log-rank tests for acquisition curves.

**Supplementary Fig. 3.**
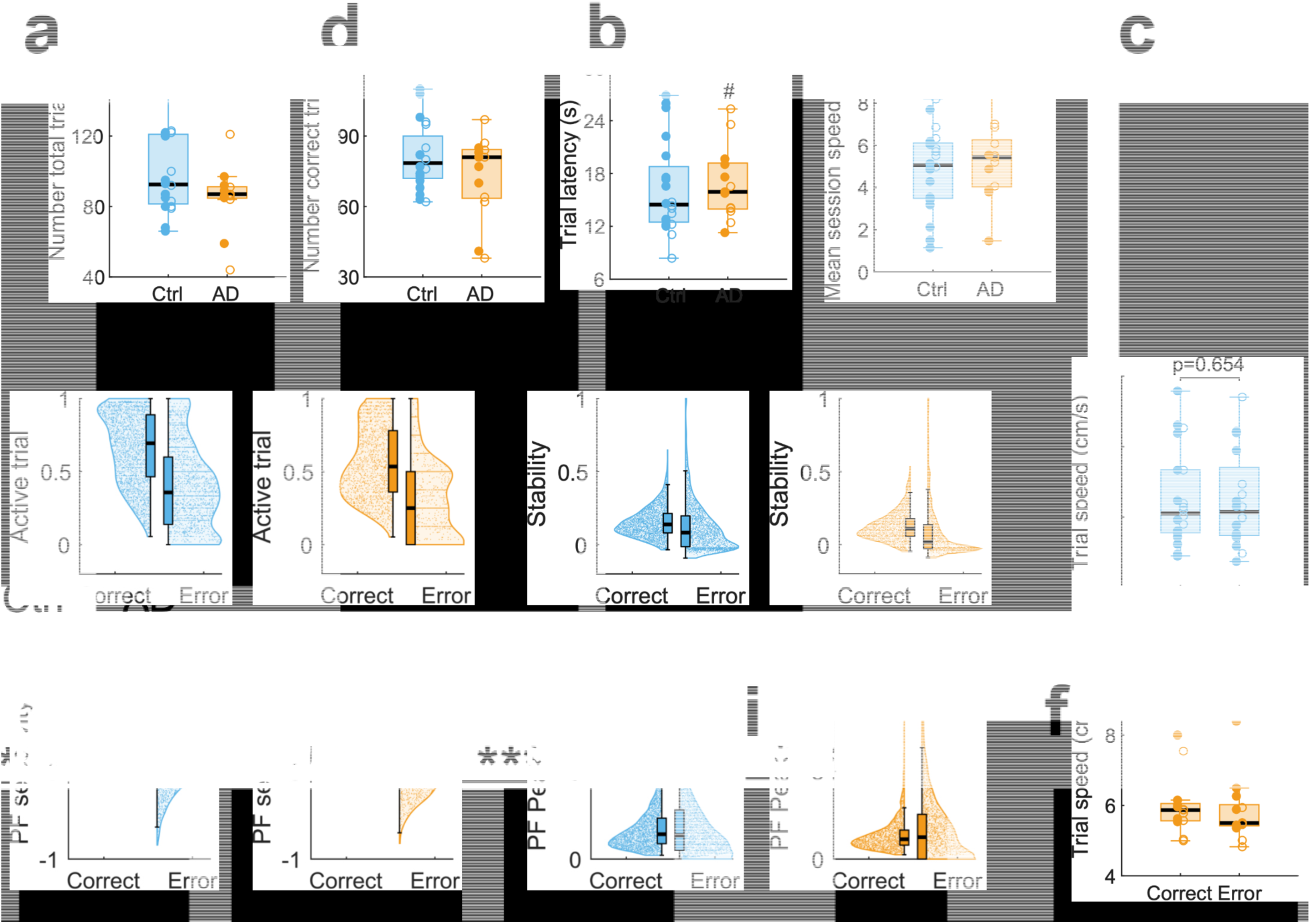
Behavioral performance and task-related place cell properties in young control and 5xFAD mice. **a-d**, Behavioral measures per imaging session in young control and 5xFAD mice: (**a**) total trial number, (**b**) correct trial number, (**c**) trial latency and (**d**) mean session speed. Each point represents one imaging session. **e-h**, Place cell activity properties during correct and error trials in control and 5xFAD mice: (**e**) active-trial ratio, (**f**) within-session stability, (**g**) place field selectivity and (**h**) peak in-field activity. Each point represents one cell. **i**, Mean trial speed per session during correct and error trials in control and 5xFAD mice. Control group includes 2952 cells from 20 sessions in 10 mice. 5xFAD group includes 1791 cells from 13 sessions in 7 mice. Box plots show medians and interquartile ranges, with whiskers indicating the data range. # indicates a genotype-by-sex interaction effect; ***P < 0.001, Statistical comparisons were performed using linear mixed-effects models for cell-level activity measures and two-sided Wilcoxon rank-sum tests for session-level behavioral measures.

**Supplementary Figure 4.**
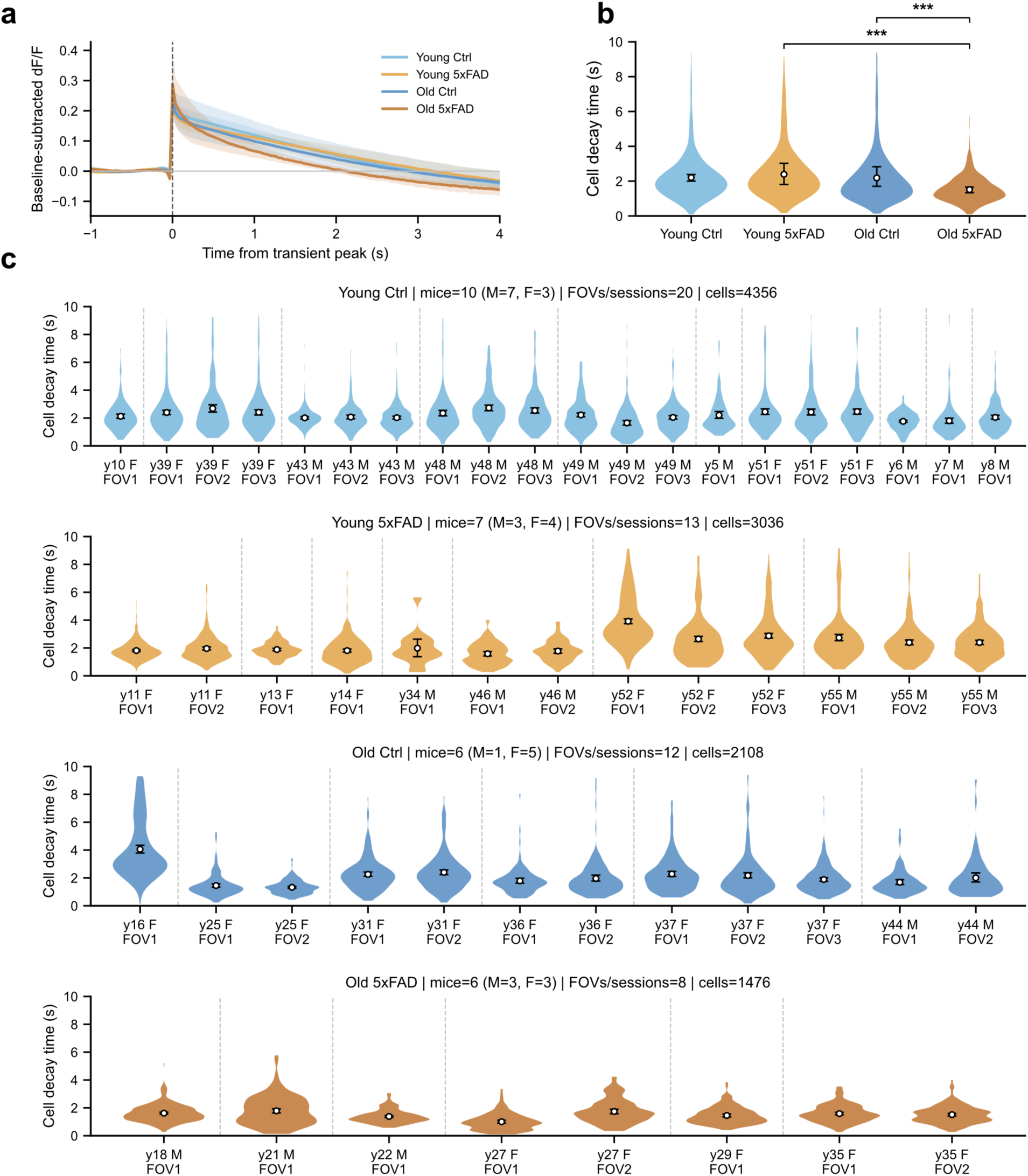
Somatic calcium transient decay kinetics across age and genotype. **a.** Average event-triggered calcium transient decay traces across the four groups. Lines show the mean baseline-subtracted dF/F aligned to the transient peak, and shaded areas indicate bootstrap confidence intervals. **b.** Distribution of single-cell decay time across the four groups. Violin plots show pooled cell level distributions for young control, young 5xFAD, old control, and old 5xFAD mice. Open circles indicate group means, and error bars indicate bootstrap confidence intervals. ***P < 0.001. **c.** Decay time distributions for individual imaging sessions. Each violin represents one FOV/session, with the corresponding mouse ID, sex, and within-mouse FOV label shown below. Dashed vertical lines separate different mice.

**Supplementary Figure 5.**
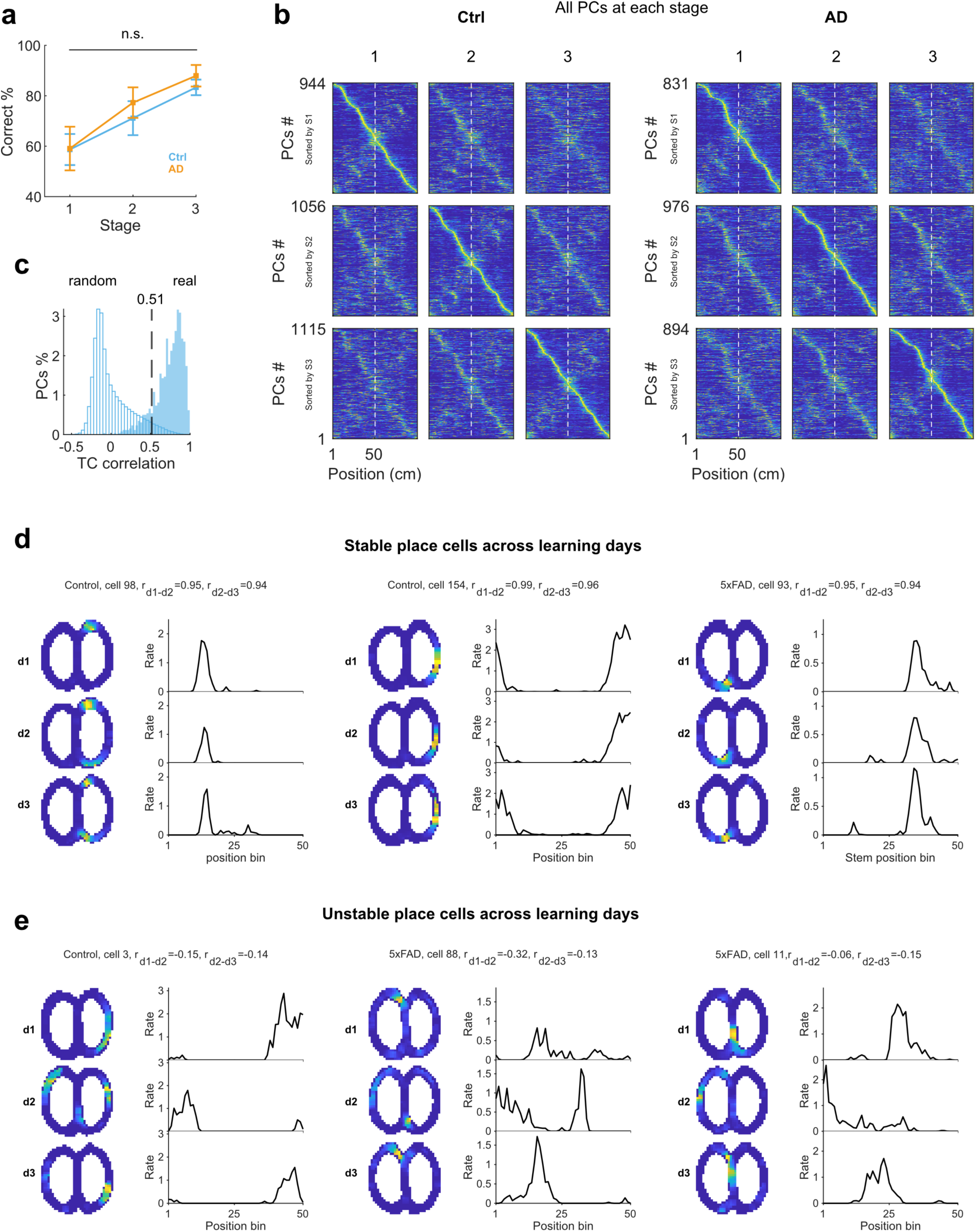
Place cells representations across training stages. **a.** Behavioral alternation performance across the three imaging stages. The gray dotted line marks the 70% performance criterion. **b.** Linearised spatial activity maps of place cells sorted according to peak position at stage 1, stage 2, and stage 3. **c.** Tuning-curve (TC) correlation threshold used to define stable place cells across stages. The observed distribution shows within-session TC correlations between spatial tuning curves averaged across odd and even trials at stage 1. The random distribution was generated from 1,000 randomly paired cells from young control mice. The threshold for cross-stage stability was set at the 95th percentile of the random distribution (>0.51) and applied to TC correlations between adjacent imaging stages. **d.** Representative stable place cells across learning stages. For each example, two-dimensional calcium activity maps and one-dimensional tuning curves are shown for stages 1-3. Adjacent-stage TC correlations are indicated above each example. **e.** Same as panel d but for unstable place cells. Data are shown as mean ± s.e.m. Control: 10 imaging sessions from 5 mice per stage. 5xFAD: 7 imaging sessions from 4 mice per stage.

**Supplementary Figure 6.**
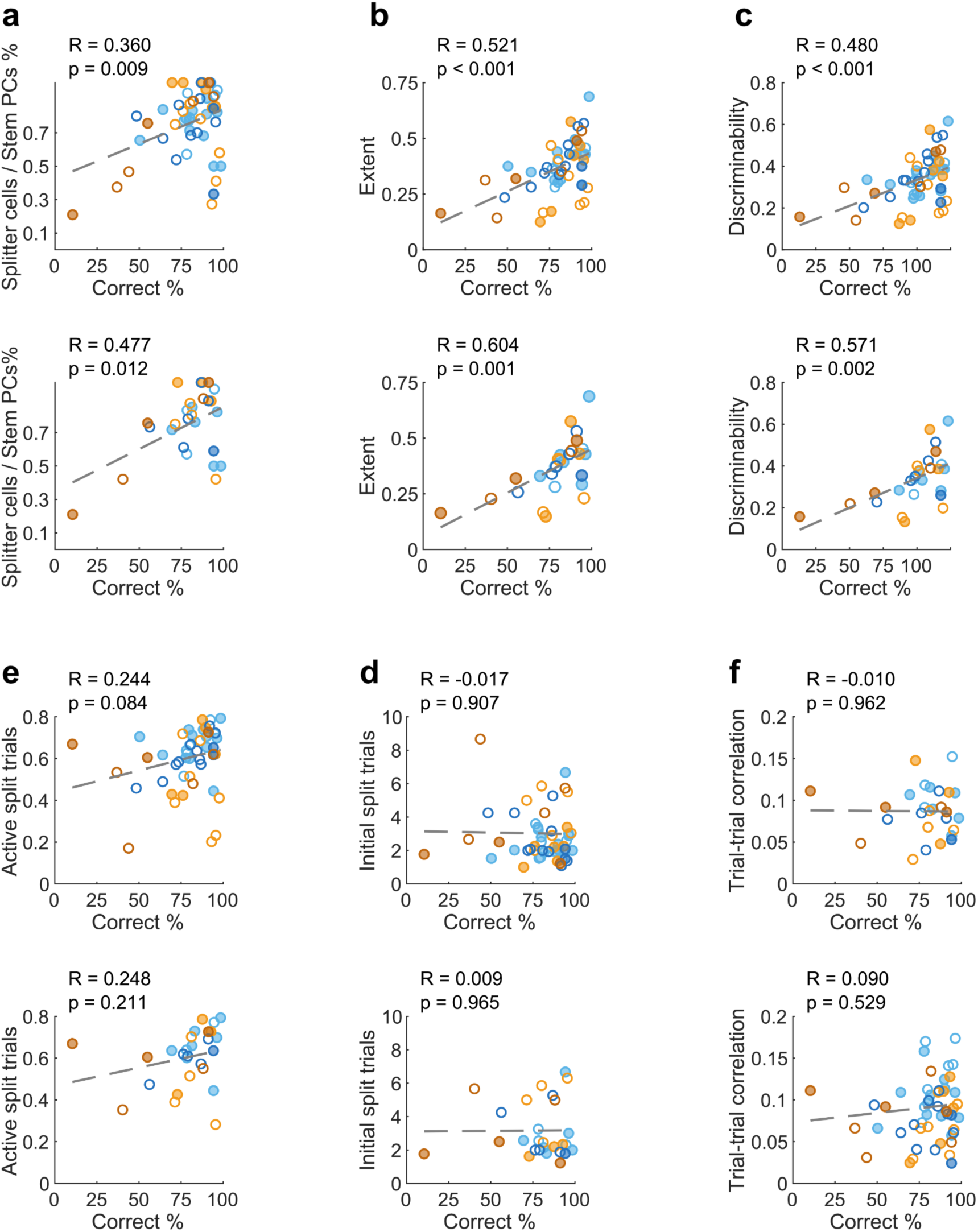
Associations between splitter cell properties and behavioral performance. Pearson correlations between splitter cell metrics and behavioral performance during the alternation task. For each metric, the upper panel shows session level values and the lower panel shows mouse level values. **a.** Proportion of splitter cells among stem place cells. **b.** Split extent, quantified as the proportion of stem positions exhibiting significant splitting activity. **c.** Split discriminability between high- and low- rate trajectories. **d.** First trial exhibiting splitting activity. **e.** Active split trial ratio. **f.** Trial to trial spatial correlation of splitter cells. Pearson correlation coefficients and P values are shown in each panel. Data were pooled across young control mice (20 sessions from 10 mice), young 5xFAD mice (13 sessions from 7 mice), older control mice (12 sessions from 6 mice), and older 5xFAD mice (8 sessions from 6 mice). Colors indicate age and genotype group. Filled and open circles indicate male and female mice, respectively.

**Supplementary Figure 7.**
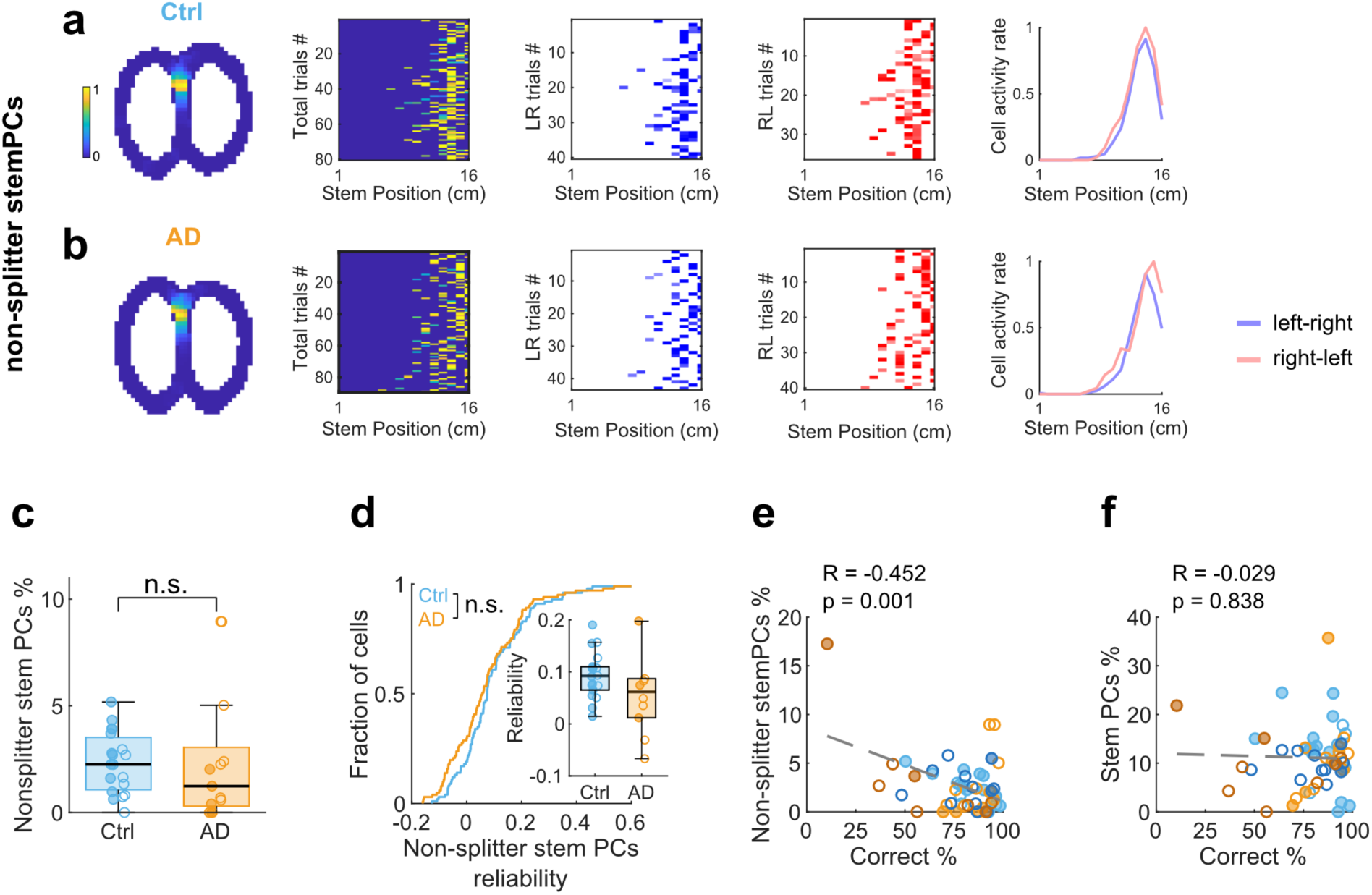
Non-splitter stem place cells. **a.** Representative non-splitter stem place cell from a control mouse. From left to right, normalized two-dimensional calcium activity map on the Phi maze from one imaging session, calcium activity across stem positions for each trial together with the corresponding linearized behavioral trajectory, calcium activity across stem positions for left-to-right trials, calcium activity across stem positions for right-to-left trials, and normalized calcium tuning curves for the two turn directions. **b.** Same as in panel **a**, but for a age matched 5xFAD mouse. **c.** Proportion of non-splitter stem place cells in control and 5xFAD mice. **d.** Trial-to-trial spatial reliability of non-splitter stem place cells. **e.** Correlation between the proportion of non-splitter stem place cells and behavioral performance. **f.** Correlation between the proportion of all stem place cells and behavioral performance. Data includes 53 imaging sessions from 29 mice. Non-splitter stem place cells included 146 cells from 32 sessions in 16 control mice and 177 cells from 21 sessions in 13 5xFAD mice. Blue and orange circles indicate control and 5xFAD mice, respectively. Filled and open circles indicate male and female mice, respectively.

**Supplementary Figure 8.**
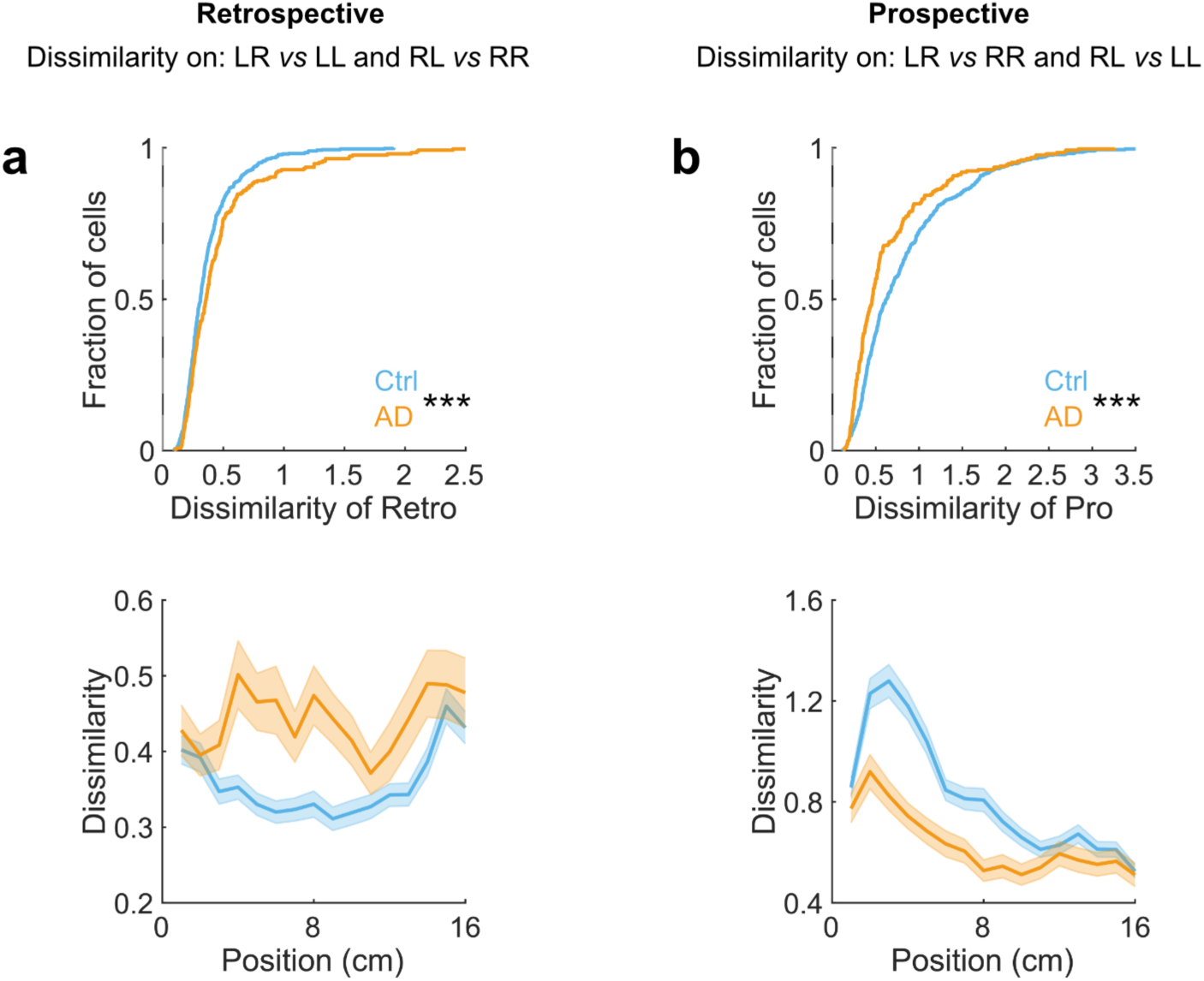
Retrospective and prospective components of trajectory-dependent coding. **a.** Top, dissimilarity values for retrospective coding, calculated from trial pairs sharing the same preceding trajectory (e.g., LR versus LL and RL versus RR). Dissimilarity was computed as −log10(P), where P was obtained from a Wilcoxon rank-sum test comparing cell activity rates between the two trial types at each spatial bin. Bottom, retrospective dissimilarity along the common stem. Lower dissimilarity indicates greater similarity between trials sharing the same preceding trajectory and therefore a stronger retrospective component. Genotype effect, F(1, 8981) = 40.55, p < 0.001; Genotype × Spatial Bin interaction, F(15, 8981) = 2.08, p = 0.008; two-way ANOVA. **b.** Same analysis as in panel a, but for prospective coding, calculated from trial pairs sharing the same upcoming trajectory (e.g., LR versus RR and RL versus LL). Lower dissimilarity indicates greater similarity between trials sharing the same upcoming trajectory and therefore a stronger prospective component. Genotype effect, F(1, 9064) = 144.46, p < 0.001; Genotype × Spatial Bin interaction, F(15, 9064) = 4.26, p < 0.001.

**Supplementary Figure 9.**
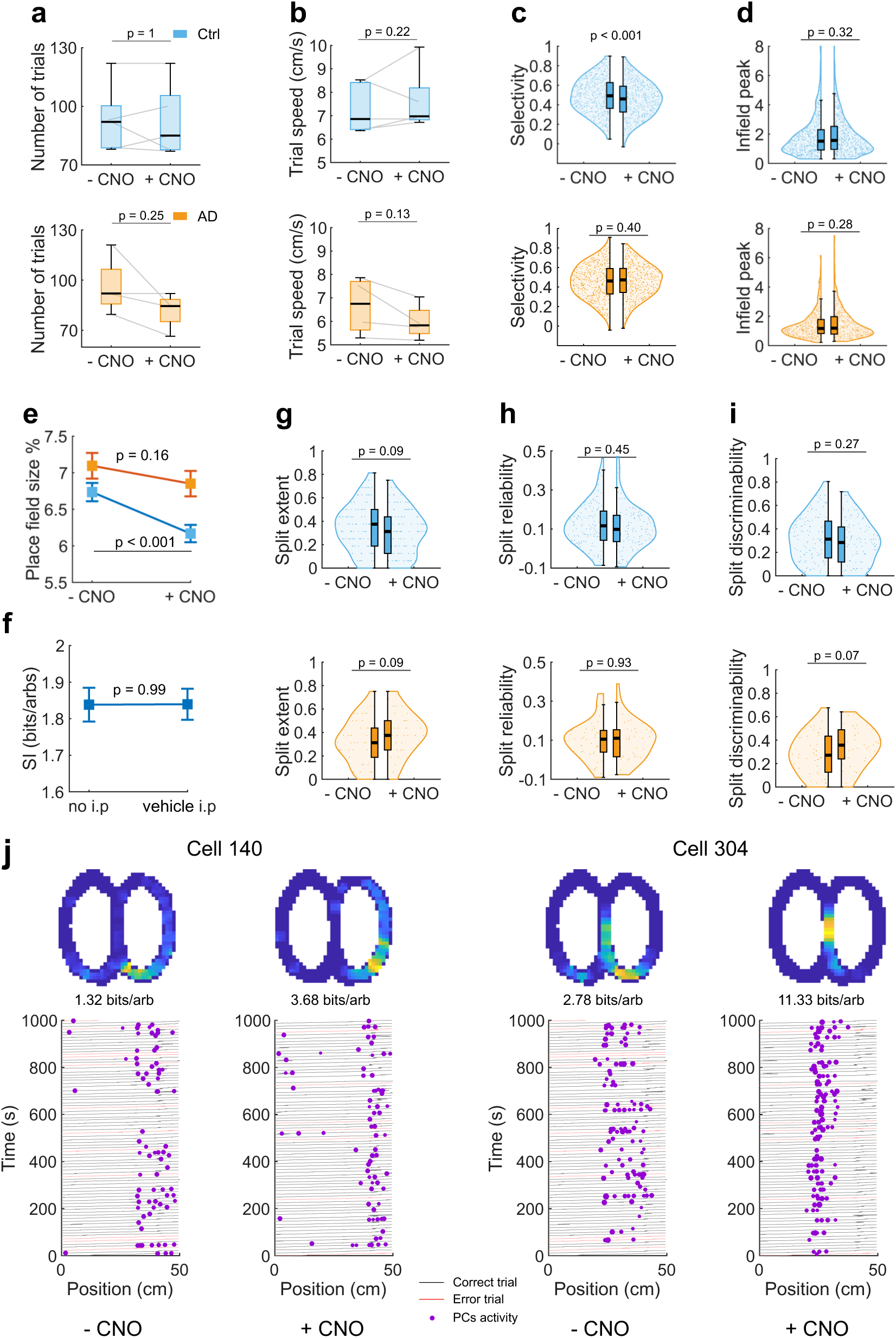
Behavioral and CA1 coding measures following medial septal cholinergic activation. **a.** Number of trials completed following vehicle or CNO administration in control and 5xFAD mice. **b.** Mean trial speed following vehicle or CNO administration. **c.** Place field selectivity of CA1 place cells following vehicle or CNO administration. **d.** Peak in-field activity of CA1 place cells following vehicle or CNO administration. **e.** Place-field size of CA1 place cells following vehicle or CNO administration. **f.** Spatial information of CA1 place cells during correct trials following no intraperitoneal (i.p.) injection or vehicle i.p. injection in hM3Dq-expressing mice. **g.** Split extent of stem place cells following vehicle or CNO administration. **h.** Trial-to-trial spatial reliability of splitter cells on the high-rate trajectory following vehicle or CNO administration. **i.** Split discriminability between high- and low-rate trajectories following vehicle or CNO administration. **j.** Representative examples of CA1 place-cell activity following vehicle and CNO administration in a 5xFAD mouse. Magenta dots indicate calcium events, gray lines indicate correct trials, and red lines indicate error trials. Cell 140 exhibited a place field on the reward arm, whereas cell 304 exhibited a stem place field with trajectory-dependent splitting activity. Data were analyzed using linear mixed-effects models with treatment (CNO versus vehicle) as a fixed effect and mouse identity as a random intercept. Ctrl, 964 detected neurons from 5 mice; 697 and 522 place cells, including 140 and 122 stem place cells, following vehicle and CNO administration, respectively. 5xFAD, 777 detected neurons from 4 mice, 522 and 449 place cells, including 46 and 41 stem place cells, following vehicle and CNO administration, respectively.

**Supplementary table 1.** The number of identified units for each mouse and each recorded FOV/session.

| Mouse ID | Genotype | Age | Group | Gender | Session /FOV ID | Total cells | Place cells | Correct trials place cells | Splitter cells |
| --- | --- | --- | --- | --- | --- | --- | --- | --- | --- |
| y5 | Ctrl | 3.7 | Young | Male | 1 | 88 | 4 | 16 | 0 |
| y6 | Ctrl | 3.7 | Young | Male | 1 | 309 | 26 | 87 | 3 |
| y7 | Ctrl | 3.7 | Young | Male | 1 | 162 | 11 | 11 | 1 |
| y8 | Ctrl | 3.7 | Young | Male | 1 | 187 | 73 | 92 | 14 |
| y10 | Ctrl | 2.96 | Young | Female | 1 | 182 | 9 | 59 | 4 |
| y39 | Ctrl | 3.06 | Young | Female | 1 | 334 | 178 | 206 | 34 |
| y39 | Ctrl | 3.06 | Young | Female | 2 | 169 | 87 | 124 | 17 |
| y39 | Ctrl | 3.06 | Young | Female | 3 | 249 | 142 | 176 | 30 |
| y43 | Ctrl | 3.06 | Young | Male | 1 | 229 | 198 | 210 | 47 |
| y43 | Ctrl | 3.06 | Young | Male | 2 | 222 | 182 | 200 | 49 |
| y43 | Ctrl | 3.06 | Young | Male | 3 | 270 | 220 | 244 | 43 |
| y48 | Ctrl | 3.6 | Young | Male | 1 | 193 | 81 | 125 | 19 |
| y48 | Ctrl | 3.6 | Young | Male | 2 | 180 | 117 | 148 | 18 |
| y48 | Ctrl | 3.6 | Young | Male | 3 | 231 | 120 | 161 | 25 |
| y49 | Ctrl | 3.6 | Young | Male | 1 | 259 | 148 | 200 | 24 |
| y49 | Ctrl | 3.6 | Young | Male | 2 | 179 | 97 | 130 | 15 |
| y49 | Ctrl | 3.6 | Young | Male | 3 | 267 | 159 | 207 | 30 |
| y51 | Ctrl | 3.6 | Young | Female | 1 | 225 | 174 | 195 | 37 |
| y51 | Ctrl | 3.6 | Young | Female | 2 | 188 | 116 | 147 | 26 |
| y51 | Ctrl | 3.6 | Young | Female | 3 | 274 | 171 | 214 | 42 |
| y11 | AD | 4.6 | Young | Female | 1 | 334 | 87 | 106 | 29 |
| y11 | AD | 4.6 | Young | Female | 2 | 266 | 102 | 155 | 29 |
| y13 | AD | 4.6 | Young | Female | 1 | 145 | 17 | 33 | 3 |
| y14 | AD | 2.5 | Young | Female | 1 | 342 | 96 | 130 | 14 |
| y34 | AD | 3.96 | Young | Male | 1 | 14 | 13 | 14 | 5 |
| y46 | AD | 2.5 | Young | Male | 1 | 99 | 20 | 59 | 4 |
| y46 | AD | 2.5 | Young | Male | 2 | 78 | 7 | 27 | 1 |
| y52 | AD | 3.6 | Young | Female | 1 | 418 | 120 | 240 | 29 |
| y52 | AD | 3.6 | Young | Female | 2 | 268 | 64 | 184 | 9 |
| y52 | AD | 3.6 | Young | Female | 3 | 369 | 85 | 256 | 23 |
| y55 | AD | 2.9 | Young | Male | 1 | 247 | 164 | 203 | 28 |
| y55 | AD | 2.9 | Young | Male | 2 | 250 | 174 | 207 | 25 |
| y55 | AD | 2.9 | Young | Male | 3 | 243 | 136 | 177 | 18 |
| y16 | Ctrl | 10.2 | Old | Female | 1 | 268 | 113 | 139 | 15 |
| y25 | Ctrl | 9.07 | Old | Female | 1 | 164 | 55 | 69 | 13 |
| y25 | Ctrl | 9.07 | Old | Female | 2 | 207 | 96 | 155 | 14 |
| y31 | Ctrl | 8.2 | Old | Female | 1 | 229 | 79 | 139 | 13 |
| y31 | Ctrl | 8.2 | Old | Female | 2 | 235 | 79 | 144 | 14 |
| y36 | Ctrl | 7.6 | Old | Female | 1 | 116 | 34 | 73 | 8 |
| y36 | Ctrl | 7.6 | Old | Female | 2 | 142 | 54 | 83 | 12 |
| y37 | Ctrl | 7.6 | Old | Female | 1 | 169 | 75 | 124 | 13 |
| y37 | Ctrl | 7.6 | Old | Female | 2 | 228 | 110 | 150 | 22 |
| y37 | Ctrl | 7.6 | Old | Female | 3 | 244 | 97 | 164 | 19 |
| y44 | Ctrl | 12.5 | Old | Male | 1 | 93 | 27 | 66 | 11 |
| y44 | Ctrl | 12.5 | Old | Male | 2 | 73 | 17 | 38 | 2 |
| y18 | AD | 8.37 | Old | Male | 1 | 284 | 106 | 109 | 13 |
| y21 | AD | 7.63 | Old | Male | 1 | 182 | 69 | 109 | 18 |
| y22 | AD | 7.63 | Old | Male | 1 | 245 | 80 | 157 | 28 |
| y27 | AD | 9.07 | Old | Female | 1 | 151 | 32 | 51 | 8 |
| y27 | AD | 9.07 | Old | Female | 2 | 109 | 29 | 55 | 11 |
| y29 | AD | 8.2 | Old | Female | 1 | 156 | 5 | 16 | 0 |
| y35 | AD | 7.6 | Old | Female | 1 | 163 | 17 | 53 | 7 |
| y35 | AD | 7.6 | Old | Female | 2 | 187 | 30 | 103 | 3 |

